# ASpediaFI: Functional interaction analysis of alternative splicing events

**DOI:** 10.1101/2020.04.12.035121

**Authors:** Doyeong Yu, Kyubin Lee, Daejin Hyung, Soo Young Cho, Charny Park

## Abstract

Alternative splicing (AS) regulates biological process governing phenotype or disease. However, it is challenging to systemically analyze global regulation of AS events, their gene interactions, and functions. Here, we introduce a novel application, ASpediaFI for identifying AS events and co-regulated gene interactions implicated in pathways. Our method establishes an interaction network including AS events, performs random walk with restart, and finally identifies a functional subnetwork containing the AS event. We validated the capability of ASpediaFI to interpret biological relevance based on three case studies. Using simulation data, we achieved higher accuracy than with other methods and detected pathway-associated AS events.

## BACKGROUND

Alternative splicing (AS) is a key regulatory mechanism that confers transcript diversity and phenotypic plasticity in eukaryotes [1]. In normal cells, splicing factors induce tissue-specific mRNA expression and embryonic stem cell differentiation [2,3]. In contrast, splice site mutations or splicing factor (SF) variants reprogram global splicing events and induce aberrant junctions in cancer cells and other diseased cells [4–6]. Aberrant AS events in cancer cells disrupt the function of tumor suppressor genes and activate the oncogenic pathways [6]. Hundreds of RNA-binding proteins, the members of the spliceosome, play a regulatory role in the cell; however, the functional effect of the spliceosome is not fully understood. As several splicing events occur simultaneously, it is challenging to infer the effects of cooperative regulation with genes and consensus pathway enrichment.

To identify the differential splicing and biological relevance, the fundamental strategy is categorized as the exon- or isoform-level approaches. The exon-level approach calculates percent spliced-in (PSI) values or total read counts from exon and junction read counts. The counts indicate exon usage, which is used for testing differential AS (DAS) events. Accurate statistical models have been developed to detect DAS that rank DAS events by significance [7–10]. However, unlike various downstream methods for gene expression analysis, the AS analysis method is restricted to inferring functional regulation induced by DAS events [11]. Previously developed application psichomics provides various downstream analyses, including the correlation between DAS and gene expression for user convenience [11].

However, they do not identify the integrative co-regulation of AS for systematically uncovering pathways. To reveal the splicing regulatory network, pCastNet identifies associations between exon and upstream regulators or downstream target genes using partial correlation. This approach requires a large number of samples (e.g. multiple tissues), and supports only the method without execution file. In spite of novel method development, exon-level approaches are restricted in DAS and their results are difficult to interpret genome-wide regulation and functions by splicing.

To uncover functional regulation, splicing studies using isoform expression also apply differential expression test and establish co-regulation network. Differentially expressed isoforms were tested like DEG test for each isoform [12]. Because isoform abundance is estimated from whole gene region, the methods result stable expression profile and DEG [12]. However these also include other limitation. Even though, major isoform differentially expressed in various conditions, isoform ratio within single gene could maintain. It is irrelevant to identify switch-like exons to regulate critical function. Nevertheless, isoform abundance is versatile to calculate expression correlations between gene pairs. To establish tissue-specific transcriptome-wide networks (TWN), previous study considered both gene and isoform expression. They identified switch-like isoforms to compute isoform ratio and established tissue-specific TWN [2,13]. TWN successfully elucidated tissue-specific molecular functions. While this method has the advantage of capturing post-transcriptional interactions, it is not adequate for tracing genomic regions of spliced exons or functional sequences like protein domains or NMD. Additionally, the isoform-level approach cannot verify cis-element usages like a donor-acceptor site, or other motifs to recognize spliceosome [6,14,15]. Therefore, a novel integrative method is required to investigate AS events and their functional interactions with partner genes as well as biological processes.

Recent studies have identified transcriptional regulation by the spliceosome in various conditions, including cancer, embryonic development, and other cellular phenotypes [3,5,6,16]. To reveal the global regulation by SFs, studies aimed at the identification of specific biological processes and the cooperative interactions were initiated [3,13,17]. Unfortunately, these approaches for identifying both splicing and associated pathways were restricted to a simple GSEA method, and independent tests for both DAS and DEG sets were performed [5,6,13]. Further, performance of multiple independent tests for splicing and gene expression does not enable the inference of global regulation by spliceosome and the interactions between AS and partner genes. Although the current gene set databases such as hallmark or REACTOME are appropriate for testing enrichment derived from gene expression [18,19], these enrichment tests using known gene sets may fail to identify novel splicing events and pertinent global interactions of the spliced genes.

Therefore, we developed a novel method ASpediaFI (Alternative Splicing Encyclopedia: Functional Interaction) to systematically identify functional AS events correlated with genes involved in pathways. We applied guilt-by-association generally used for gene expression analysis to splicing regulation. In order to reveal transcriptome-wide global regulation of both spliced genes and non-spliced genes, we established a heterogeneous interaction network for both genes and AS events. To increase interpretation availability, pathways including gene set information were also included to the network. Our applications explore splicing subnetwork regulated by SF conditions using discriminative random walks with restart (DRaWR). The algorithm has been applied to various heterogeneous networks like gene co-expressed interactions, sequence homology, or transcription factor-binding motif [20,21].

Random walk with restart (RWR) algorithm explores the interaction networks from a query gene set - called seed, and finally ranks nodes based on association with the query. To confirm whether our analysis method produces a biologically relevant result, we applied our method to three RNA-Seq datasets, which included samples from cancer patients with the SF variant and SF knockdown cells. We compared our results for three RNA-Seq dataset with previous results and other tools. The result was verified in various aspects like AS event types’ proportion, biological relevance, discriminative power, and other parameters. We also evaluated the performance of our method using simulated dataset. ASpediaFI is available in Bioconductor (https://bioconductor.org/packages/ASpediaFI).

## RESULTS

### ASpediaFI algorithm and analysis workflow

ASpediaFI identified a subnetwork from a heterogeneous network established using gene-gene interactions, containing gene-AS and gene-pathway interactions. The interaction network was based on the concept of guilt by association, which states that associated or interacting genes are more probable to share function [21]. We expanded the network by adding AS events and pathways to the feature nodes. Quantitative information weighting network edges were collected from PSI, gene expression and pathway gene sets. The ASpediaFI workflow starts with data preparation and sequentially follows through heterogeneous network establishment, subnetwork exploration, and further downstream analysis. During the data preparation step, our method identifies AS events from gene model annotation, collects gene expression, calculates PSI profile, and refers pathway gene sets and gene interaction data collected from public databases (Figure 1A). ASpediaFI integrates the processed data to construct a heterogeneous network that contains gene nodes and its feature nodes representing AS event and pathway. Before executing the algorithm, the adjacency network is normalized within the feature nodes and for all nodes. Next, to explore the subnetworks, our method performs DRaWR on the heterogeneous network using previously defined relevant query gene sets collected from DEGs (Figure 1B, blue node) [21]. In the first stage, our algorithm explores the highly ranked feature nodes from the query set. We then extract a subnetwork from these feature nodes chosen from the first stage and all gene nodes, including associated edges (Figure 1B). Next, ASpediaFI performs second stage RWR for gene nodes to rank again and additionally calculates *P*-values by permutation tests to eliminate background effects like query gene size. For user convenience, our tool provides further analyses, including GSEA and data visualization. More details of our algorithm are described in the Method section.

**Figure 1.**
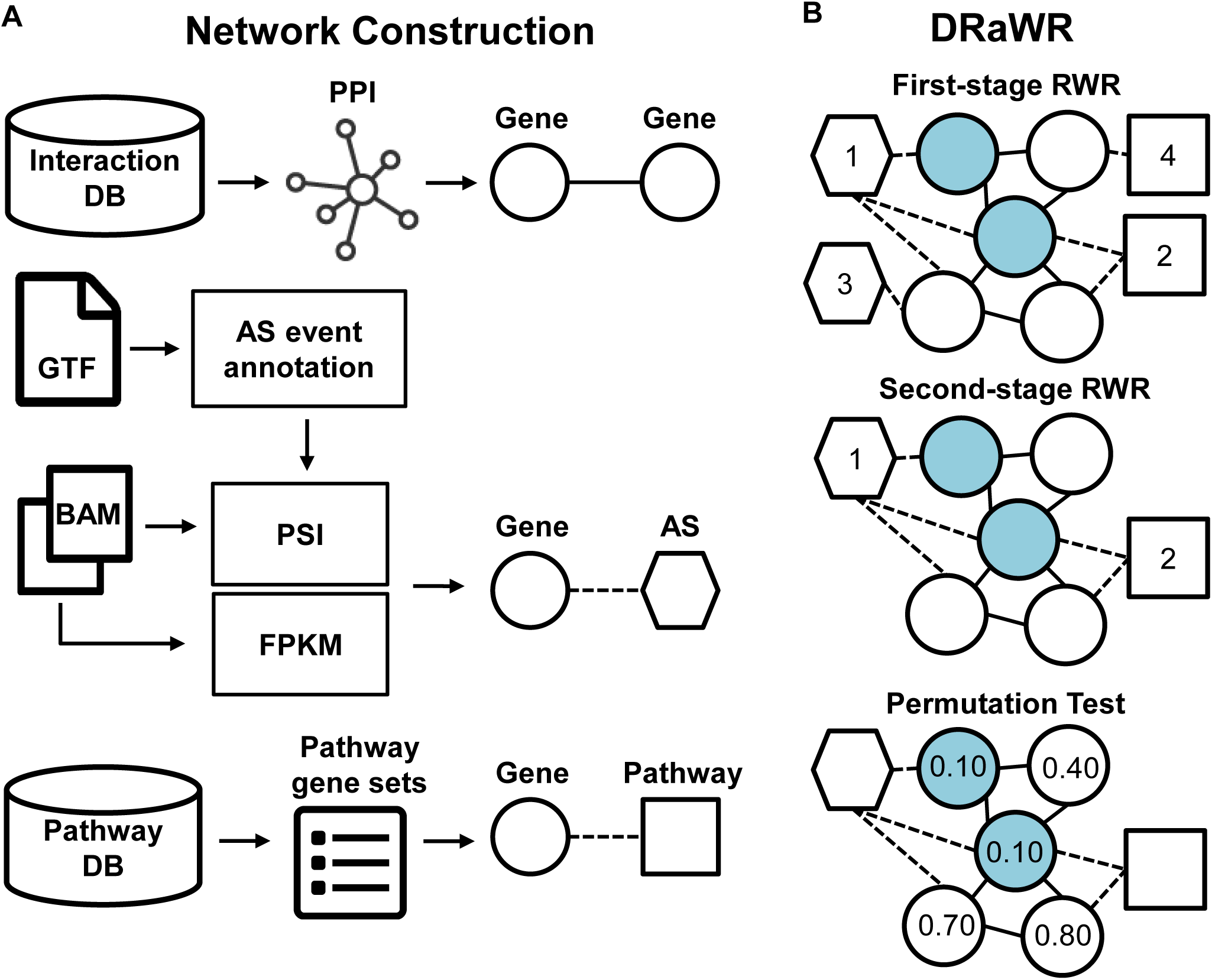
ASpediaFI workflow and DRaWR algorithm to identify AS interaction subnetwork. A) ASpediaFI establishes a heterogeneous network using gene interaction, gene-AS correlation, and gene-pathway association data. AS events are annotated from a GTF file, and PSI calculation using BAM files is also embedded. Public gene sets are referred for gene-pathway associations. B) A heterogeneous network is composed of genes and its feature nodes, AS events and pathways. Gene-gene and gene-AS interaction edges are weighted by correlations of gene expression and PSI values. Next, all edge weights are normalized for each type of feature interaction and each column. The first stage RWR explores a heterogeneous network starting from nodes in a query gene set (blue nodes). The second stage RWR finalizes scores within a query-specific subnetwork derived from the first stage. ASpediaFI additionally computes permutation *P*-values of the gene nodes to eliminate the effect of the background gene set.

### Alternative splicing analysis using three RNA-Seq datasets applying ASpediaFI

To verify the capability of ASpediaFI, we analyzed three RNA-Seq datasets representing the following: myelodysplastic syndrome (MDS), stomach cancer (STAD), and RBFOX1-knockdown cell lines. MDS and STAD were collected from cancer patients, and RBFOX1 has replicated samples of a relatively smaller size (n = 5 per condition). The datasets contain SF mutations or down-regulations. We compared the SF deficiency profiles with the wild-type using ASpediaFI and investigated whether our DAS sets determine the splicing pattern and cis-element usage by the spliceosome. The biological relevance of our highly ranked pathway result was delineated by referring to previous studies, and the consistency of GSEA in using gene expression was also evaluated. Additionally, we tested how much our DAS set was enriched in known and novel pathways or how much our result was coherent based on other known AS signatures compared to other methods. In a further overall investigation of the DAS set, we thoroughly examined the DAS events belonging to known and novel pathways compared to other results. Each spliced gene was reviewed for biological relevance and functional consistency with identified pathways. Additionally, functional sequence features like protein domain and NMD that exist on spliced exons were extensively investigated and compared with other results [22].

### Case study 1: Three splicing factor mutations in myelodysplastic syndrome induce the dysregulation of heme metabolism

We investigated AS events in RNA-Seq samples from MDS patients (n = 84) with SF deficiency on SF3B1 (n = 28), SRSF2 (n = 8), and U2AF1 (n = 6) using three respective query gene sets of 112, 107, and 96 differentially expressed genes [13]. By comparing SF mutations (MUT) with wild-type (WT) samples, we identified 281, 269, and 285 AS events and 19, 31, and 15 pathways, respectively, for SF3B1, SRSF2, and U2AF1 (Additional File 1: Table S1). Proportions of each AS event type are summarized in Figure 2.A. RI (37.9□53.7%) was most frequently detected in three cases, and the frequency of A3 events was next (Additional File 2: Table S2). The dominant occurrence of RI and A3 events in our result is consistent with a previous MDS analysis study using rMATS [4,6,13,23–26]. However, in the previous study, a more refined final DAS set from two comparisons using both WT and healthy control samples as controls, were selected [7,23]. When considering a single comparison with WT samples in rMATS as we did, SE showed the largest proportion across the three SF analyses (34.1 ∼ 59.4%; Additional File 2: Table S2). When comparing with results from the other two methods (performed in the final section of evaluation), SUPPA2 detected A3 (40.3 %) most frequently, followed by SE (28.7 %) and RI (19.4 %) [8]. MISO showed a similar pattern to that of rMATS (SE 25.5 %, A3 18.4 %, and RI 25.7 %) [9]. Three SFs, SF3B1, SRSF2, and U2AF1, are known to recognize the 3′ splice sites (acceptor sites) [15]. Therefore, our method minimized bias toward specific AS event types and reflected the role of spliceosome recognizing cis-elements. SUPPA2 was also able to project the characteristics of the spliceosome.

**Figure 2.**
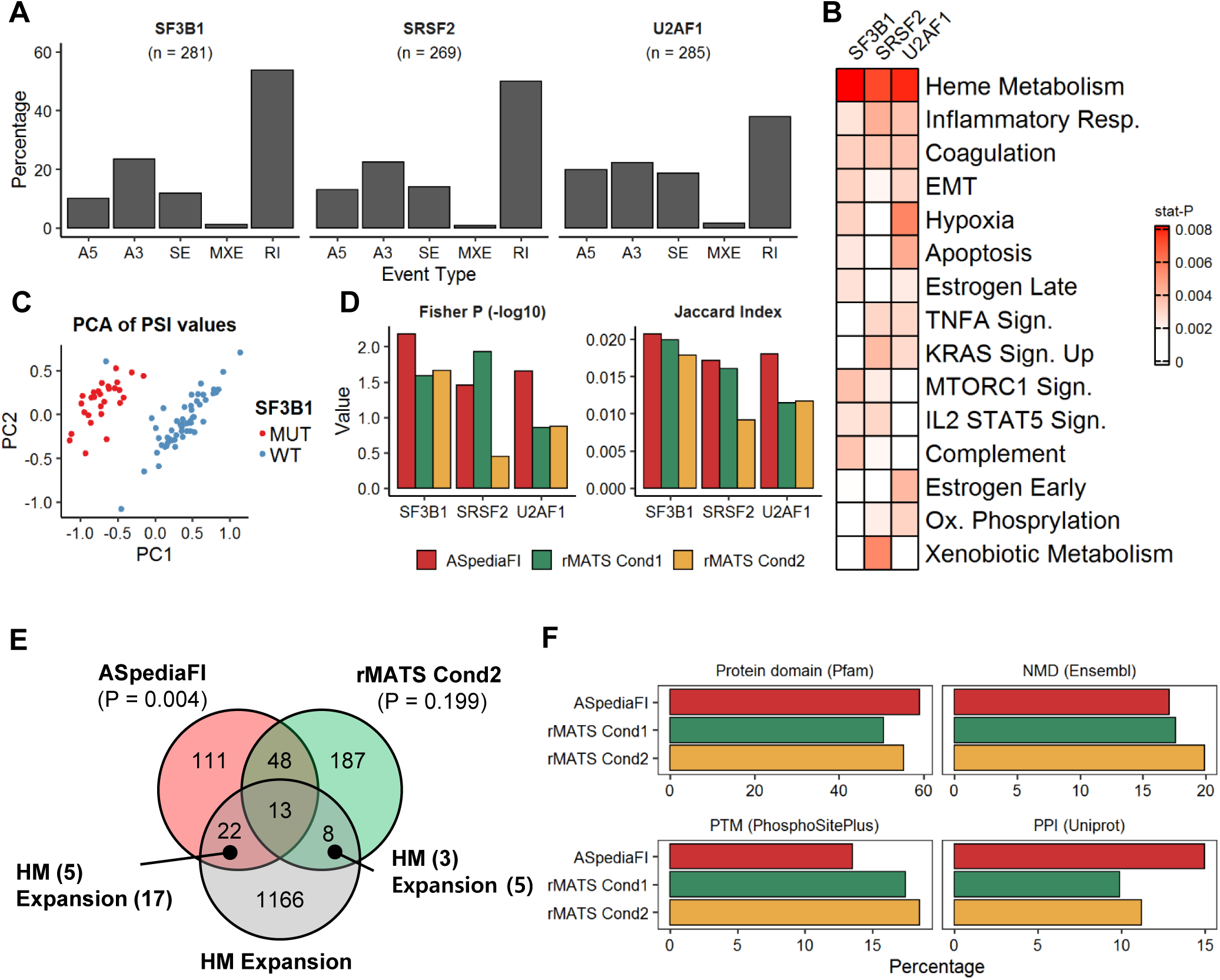
MDS patient RNA-Seq dataset analysis to identify AS events and pathways regulated by SF3B1, SRSF2, and U2AF1 mutations. A) Percentages of five AS types identified by ASpediaFI for three SF MUT cases. B) Heatmap of the top 15 pathways ranked by stat-Ps. C) PCA plot derived from PSI profiles of SF3B1 MUT-associated 281 events. PC1 (x-axis) and PC2 (y-axis) indicate principal component 1 and 2. D) Two barplots of AS event enrichment comparison in HM pathway gene set for three conditions: ASpediaFI, rMATS Cond1, and Cond2. One is negative log-scale *P*-values of Fisher’s exact method and other’s Jaccard indices. E) A Venn diagram of genes related to SF3B1-associated AS events identified by ASpediaFI and rMATS (Cond2) compared with the HM expansion set containing both HM pathway gene set and interacting novel gene set. For testing enrichment with HM expansion, *P*-values for ASpediaFI and rMATS were calculated by Fisher’s exact test. In two exclusive intersections of ASpediaFI (n=22) and rMATS (n=8), ASpediaFI detected more events (n=10) in the expansion set than rMATS (n=5) as well as total events in two exclusive intersections. F) Percentage barplots of AS events to contain four functional sequence features, protein domain, NMD, PTM, and PPI. It was also compared with rMATS Cond1 and Cond2.

To delineate pathway regulation for each SF MUT, we presented hallmark pathways highly ranked by stat-P (Figure 2B; further details of stat-P described in the Method section). The heme metabolism (HM) pathway was top-ranked in all three analyses. Coagulation, hypoxia, oxidative phosphorylation, inflammatory response, and estrogen receptor signal pathways were also revealed to be regulated by the three SF MUTs. The previous MDS study used a commercial software, IPA for pathway analysis, which ranked sirtuin signaling as the first and heme biosynthesis as the second [13]. As the sirtuin pathway was absent in the hallmark pathway set, our result is remarkably similar to that of the previous study.

For additional validation, we evaluated the discriminative power of our AS event set and compared the enrichment to biological function with the rMATS result. Specifically, we investigated the result of the SF3B1 analysis. As shown in a scatterplot of principal component analysis (PCA), the PSI profile of 281 AS events accurately discriminated between the MUT and WT samples (sensitivity: 100%, specificity: 96.3%; Figure 2C). To compare and to evaluate pathway enrichment of our AS event set, we executed rMATS and obtained DAS sets for two conditions, one with the same criteria adopted in the previous MDS study (Cond1; n =596) and another with a more strict option (Cond2; n = 367) [13]. Previous literature in combination with our findings (Figure 2B) suggests that hematopoietic malignancy, HM, and heme biosynthesis are dysregulated by SF3B1 mutation in MDS or U2AF1 in other blood cancers [4,6,13,23–26]. Therefore we selected the HM pathway as a true gene set. ASpediaFI demonstrated the best overall performance from the perspective of both Fisher’s exact test *P*-value and Jaccard index in all three SF MUT analyses except SRSF2, where rMATS Cond1 showed the lowest *P*-value (Figure 2D). We additionally examined our AS event genes and their specific functions using the Venn diagram to compare the three sets from ASpediaFI, rMATS Cond2, and HM expansion set (Figure 2E). We chose rMATS Cond2, which performed better in the SF3B1 analysis over Cond1. We generated the HM expansion set by merging the HM pathway gene set with interacting genes in our PPI network in order to investigate novel candidates for genes regulating the pathway by alternative splicing (more details are described in method). AS event genes of ASpediaFI (Fisher’s exact test *P*-value = 0.004) are more significantly enriched in the HM expansion set than in rMATS Cond2 (*P*-value = 0.199). Meanwhile, we explored several functional sequence features involved in splicing regions using the ASpedia database. The AS events generated by our analysis were involved in more protein domains, nonsense mediated-decays (NMD), and isoform-specific protein-protein interactions (PPI) than those generated by rMATS Cond1 and Cond2 but contained fewer post-translational modifications (PTM) and repeat regions (Additional File 2: Table S4; Figure 2F) [22].

We examined the biological function of spliced genes in two distinct mutually exclusive sets of ASpediaFI (n = 22) and rMATS (n = 8) overlapping with the HM expansion set (Additional File 2: Table S3). We divided the HM expansion set into two, known genes that belong to the HM pathway and novel genes adjacent to the genes in the HM set. Total events in the exclusive sets were detected at a higher frequency with ASpediaFI. Novel AS genes were also detected more efficiently with ASpediaFI (n=17; rMATS n=8). We identified two known splicing genes NARF and SNCA, that are directly associated with MDS belonging to the HM pathway (Additional File 2: Table S3). Interestingly, only ASpediaFI detected an AS event on the synuclein alpha (*SNCA*) gene, and the ASpedia database identified the ‘synuclein’ domain in the AS inclusion region (Additional File 2: Table S3), which has been shown to interact with sirtuin 2 [27]. As we already described in previous, sirtuin signaling was not able to detect in our result (Figure 2B). However we successfully identified SE event of the sirtuin signal-associated gene, *SNCA* and our result implies to engage spliced genes in both heme metabolism and sirtuin-1 autophagy pathway like previous finding [28]. We also exclusively identified a RI event of NARF in the C-terminal’ domain of the ‘Iron only hydrogenase large subunit’ where the event induces Alu-exon insertion and affects substrate-binding affinity or catalytic activity in MDS [25]. The ASpediaFI results also included more novel AS events (HM expansion set) (47% of 17) involved in protein domains compared to rMATS (40% of 5; Additional File 2: Table S3). Among the novel events identified by ASpediaFI, we found a RI event of CDC37. The gene is regulated by Hsp90 during the biogenesis of the active conformation of the heme-regulated eIF2α kinase, and spliced site is critical to the loss of the ‘Hsp90 binding’ domain [29]. In summary, ASpediaFI identified more number of novel AS events than rMATS. These findings can be interpreted as evidence that ASpediaFI efficiently detects novel and functionally important AS events.

### Case study 2: EMT pathway in stomach cancer induced by ESRP1 and the representative AS events

Epithelial regulatory splicing factor, ESRP1, is down-regulated during epithelial-mesenchymal transition (EMT) and plays a critical role in tumor progression [30,31]. We performed analysis on the TCGA STAD RNA-Seq dataset to examine ESRP1-related AS events, associated pathway regulation. Additionally, we investigated the consistency of our results by comparing it with GSEA using gene expression to verify our method by performing integrative analysis. Samples were classified into ESRP1 high (n = 41) and low (n = 42) groups based on ESRP1 mRNA expression (RPKM). ASpediaFI identified seven pathways and 293 AS events (Additional File 1:Table S1). The PSI profile of the detected AS events provided a powerful discriminatory performance (Sensitivity: 100%, Specificity: 69%; Figure 3A). The proportions of five AS event types are presented in Figure 3B. SE was identified in 66%, and it was three times the sum of (22%) of A3 and A5. In additional DAS analysis using SUPPA2, SE events (57%) were detected most frequently. Percentages of five AS types identified by ASpediaFI consistently resembled those detected by SUPPA2, as already uncovered in case study 1. In pathway analysis, ASpediaFI ranked the EMT pathway on top and consequently identified EMT-associated pathways such as ‘myogenesis’ and ‘apical junction’ (Figure 3C). To compare gene expression-based analysis with ours, we estimated pathway scores for each sample using GSVA from the gene expression profile and compared them with our pathway rankings (Figure 3C) [32]. The GSVA result resembled our rankings except for two pathways, ‘IL2-STAT5 signaling’ and ‘UV response down,’ which exhibited lower relevance than EMT and myogenesis.

**Figure 3.**
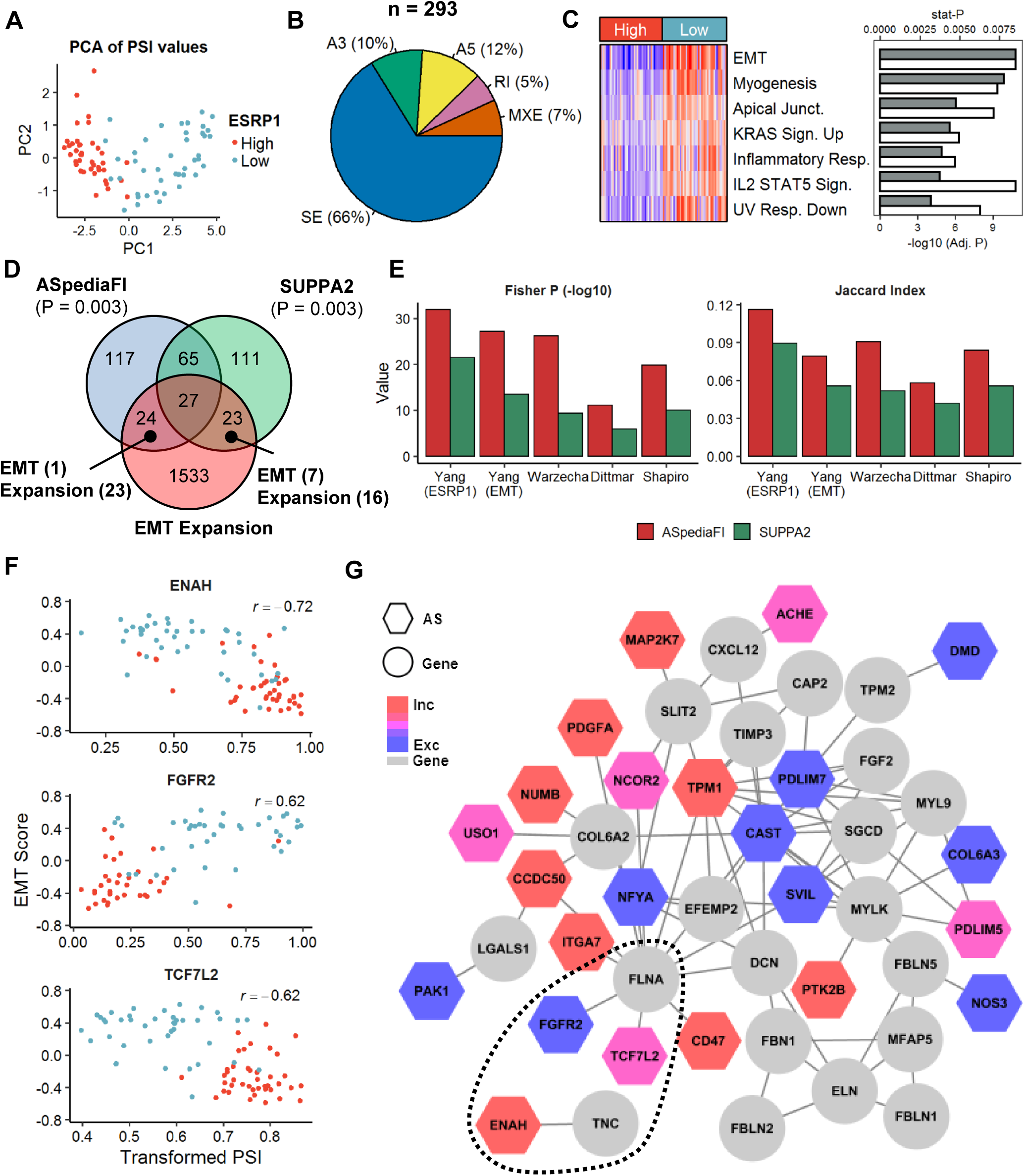
AS events associated with the EMT pathway regulated by splicing factor ESRP1. A) PCA scatter plot using PSI profiles using 293 AS events. B) Percentage pie chart of five AS types. C) Pathway identification comparison between our method and gene expression-based analysis. Seven pathways in heatmap row were chosen from ASpediaFI pathway ranking, and columns were ordered by high and low groups. The heatmap demonstrates pathway-level GSVA scores estimated using gene expression profiles. The barplot on the right layout presents both our stat-P values (gray) for pathway ranking and log-scaled adjusted *P*-values (white) of GSVA scores comparing between ESRP1 high and low groups. D) Venn diagram of ASpediaFI, SUPPA2, and the EMT expansion gene set. *P*-values for two AS sets denote enrichment with EMT expansion set. E) Status barplots to investigate AS event consistency identified by ASpediaFI and SUPPA2. Five EMT splicing gene signatures (Yang ESRP1 [33], Yang EMT [33], Warzecha [34], Dittmar [35], and Shapiro [31]) were collected, and Fisher’s exact test *P*-values and Jaccard indices were calculated. F) Scatter plots between EMT pathway scores (y-axis) by GSVA and square-root-transformed AS event PSI values (x-axis) for three AS events, ENAH, FGFR2, and TCG7L2. Correlation coefficients were added to each plot. Blue dots indicate low group and red dots indicate high group. G) A gene-AS interaction subnetwork identified by ASpediaFI. Circle nodes denote gene nodes, and hexagons are AS events. AS event nodes were filled in color by dPSI values. To extract smaller size EMT-relevant subnetwork for generating plot, we eliminated gene nodes belonging to the EMT expansion set with log2 fold change < 2 and AS nodes of | dPSI | < 0.25. Multiple edges of one AS node were trimmed except the one with the maximum score. The dotted line ellipse indicates the interactions of three spliced genes (Figure 3F).

To investigate the biological function and novelty of spliced genes, we compared two AS gene sets inferred from ASpediaFI and SUPPA2 with the EMT expansion set (Figure 3D). The two gene sets were equivalently enriched in the expansion set (Fisher’s exact test *P*-value < 0.003). When retrieving functional sequences of DAS events from ASpediaFI and SUPPA2 (Additional File 2: Table S4), AS events were comparably enriched in protein domains for ASpediaFI (32.5%) and SUPPA2 (33.1%). The frequency of NMD was slightly higher in ASpediaFI, and SUPPA2 was better at identifying repeat regions. PTM and PPI were remarkably much more frequently identified by ASpediaFI (37.0%, 35.8%) than SUPPA2 (29.9%, 24.0%). Meanwhile, ASpediaFI exclusively identified more novel AS events (n = 23) than SUPPA2 (n = 16). Moreover, the novel events identified by ASpediaFI were more involved in protein domains (ASpediaFI: 34.8 % of 23 events, SUPPA2: 25% of 16; Additional File 2: Table S5). On comparing the DAS sets from ASpediaFI and SUPPA2 with five known EMT or ESRP1-associated splicing signatures [31,33–35], the results of the Fisher’s test and Jaccard indices were notably better for the ASpediaFI DAS set across all signatures (Figure 3E).

Notably, our result identified novel events, ENAH SE, FGFR2 MXE, and TCF7L2 SE, which were neither present in the hallmark EMT pathway gene set nor detected by SUPPA2. The three events were also identified in all five splicing signatures (Figure 3E). PSI values of these splicing events exhibit strong correlation coefficients (| *r* | = 0.62 ∼ 0.72) with EMT pathway scores calculated by GSVA based on gene expression (Figure 3F). Our three events were also present in the EMT-associated submodule extracted by applying stringent cutoffs (gene log2 fold change > 2 and AS | dPSI | > 0.25) (Figure 3G). Our network revealed the functional interactions of TCF7L2 and FGFR2 with FLNA to be a network hub and to regulate EMT in tumor cells [36]. Occurrence of the representative three AS events in the genomic regions lead to changes in the protein domain, and these were shown to be strongly involved in EMT-associated functions based on previous literature [16,37,38]. ENAH, an actin cytoskeleton regulatory gene, is spliced, and exon11a is skipped on the EVH2 domain (Additional File 3: Figure S1). FGFR2 MXE generates two isoforms: FGFR2-IIIb, which is exclusive to epithelial cells and FGFR2-IIIc, which causes a switch from the mesenchymal isoform and induces a change in ligand binding specificity, thereby regulating cell proliferation and differentiation (Additional File 3: Figure S1) [16]. TCF7L2 SE is present in the ‘N-terminal CTNNB1 binding’ region, FGFR MXE in the ‘Immunoglobulin I-set domain,’ and ENAH in the ‘EVH2 domain’ (Additional File 3: Figure S1). TCF7L2 SE in the CTNNB1 binding domain has an impact on the activity of Wnt/β-catenin target genes, and its deficiency was verified as the depletion of a proliferative cell compartment in the intestinal epithelium in mouse [38]. Its switch-like exon usage was revealed to be associated with invasive and mesenchymal-like breast tumors [37].

### Case 3: Splicing events uncover neuronal development by RBFOX1 knockdown

AS events mediated by RNA-binding protein RBFOX1 regulate neuronal development and pertain to brain diseases like autism [5,14]. We analyzed the RBFOX1 knockdown RNA-Seq dataset of primary human neural progenitor cells, which included five RBFOX1 knockdown samples and five control samples. In order to be consistent with the previous study, we changed the reference pathway gene set to GO level 5 [5]. Finally, ASpediaFI identified 291 AS events and nine pathways (Additional File 1: Table S1). A3, RI, and SE were frequently detected, and MXE was the least predominant (Figure 4A). To verify the result, our AS genes were compared with three relevant gene signatures (autism, RBFOX1, and RBFOX2) and three controls (mitrochondrial, ataxia, and epilepsy) obtained from the previous study using the Jaccard index (Figure 4B) [5]. Relevant signatures were collected from spliced gene analysis results of autism (n = 247), RBFOX1 (n = 1103), and RBFOX2 (n= 1681). Controls were randomly selected from known gene sets, mitochondrial (n=310), ataxia (n=51), and epilepsy (n=46). Relevant signatures exhibited higher similarity to our AS gene set in terms of the Jaccard index compared to that of the control set (Figure 4B). In accordance to the pathway ranking of our analysis, neurogenesis, neuron differentiation, and nervous system development pathways were induced in response to RBFOX1 knockdown (Additional File 1: Table S1).

**Figure 4.**
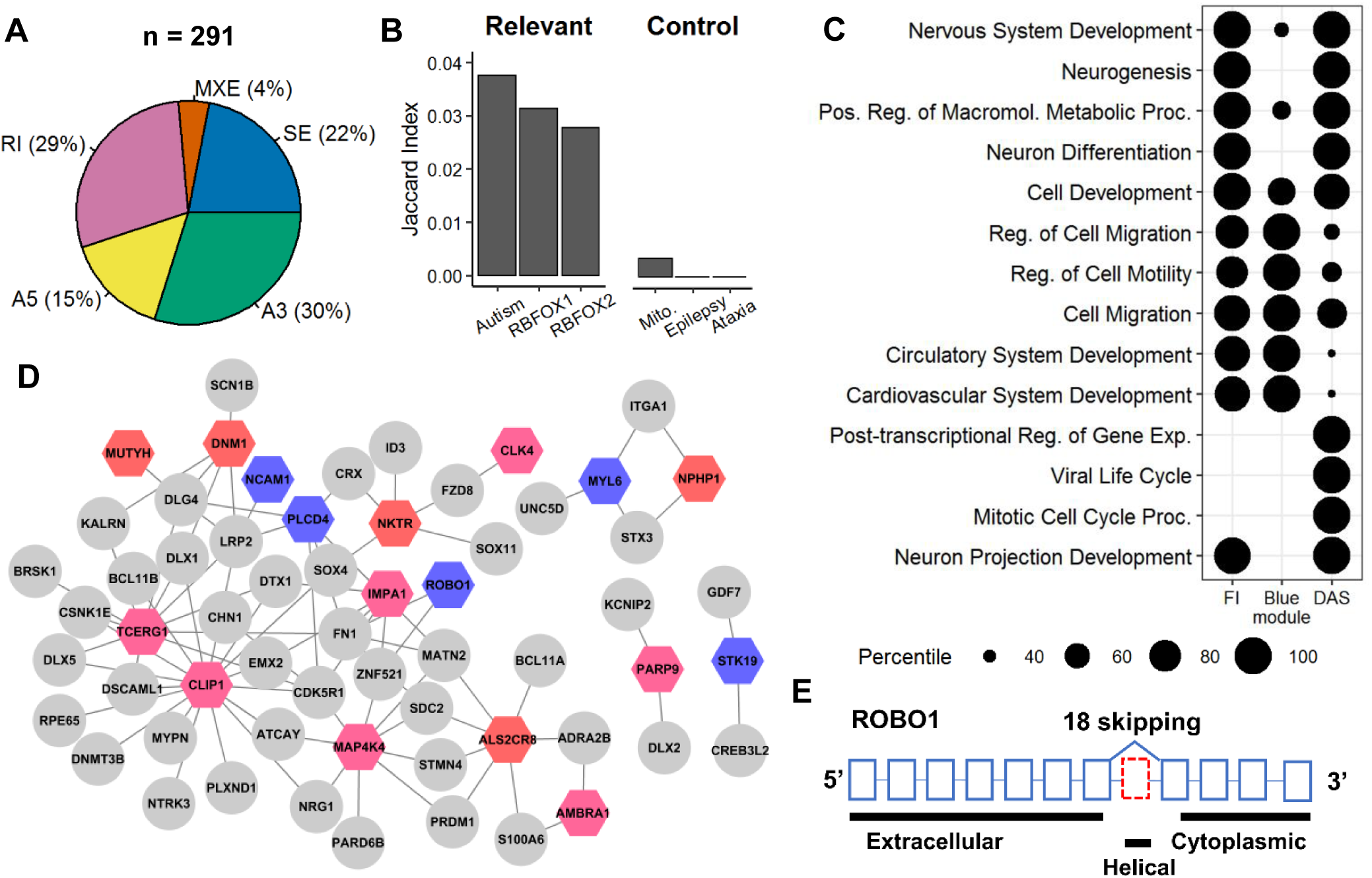
Analysis of the RBFOX1 knockdown RNA-Seq dataset. A) A pie chart showing the proportion of 5 AS event types. B) Jaccard index barplots between our result and splicing gene signatures collected from a previous study [5]. Three relevant RBFOX1-associated splicing gene sets were overlapped with three controls. C) Dot plot for percentile ranks of GO terms (row) from gene sets (column) by three different methods, our genes extracted by permutation *P*-values (FI), neuronal development genes identified by WGCNA referring gene expression (Blue Module), and differentially spliced genes (DAS). The last two gene sets were derived from the previous study result. D) RBFOX1-associated subnetwork that ASpediaFI identified. To extract a smaller size subnetwork, we eliminated gene nodes belonging to neuron differentiation set with log2 fold change < 0.25 and AS nodes of | dPSI | (< 0.15). E) Exonic structure of exon 18 skipping (red) and protein domains of ROBO1.

To evaluate the pathway detection performance, we compared our results with those of the previous study [5]. The study generated two sets of SE events with differential exon inclusion and exclusion. The biological process was also investigated by GSEA for each AS set. In further GSEA using the AS event set, the previous study identified a subnetwork regulated at the gene expression level. We combined the two AS sets into one ‘DAS’ set and used a co-expressed subnetwork gene set named ‘Blue module’ from the previous study [5]. To verify the gene set enrichment in biological process detection potential, these two gene sets were compared with the highly scored genes (permutation *P*-value < 0.05) identified by our method. We chose the top five GO terms from the GSEA result of the three gene sets and computed their percentile ranks (Figure 4C). The ‘Blue module’ was enriched in cell migration and motility but failed to detect neuronal differentiation and neurogenesis. In contrast, we observed that nervous system development was more enriched than cell migration and motility in the ‘DAS’ gene set (Figure 4C). Unlike these two signatures, ASpediaFI successfully identified the most relevant biological processes associated with neuronal development on top percentile rank GO terms except for post-transcriptional regulation, viral life cycle, and mitotic cell cycle (Figure 4C, the first column FI). This result illustrates the advantage of our integrative approach based on both gene expression and PSI profiles and the limitation of independent gene set tests (Blue module and DAS) for analyzing splicing-associated biological functions.

We identified an RBFOX1-associated module within the heterogeneous network (Figure 4D). The subnetwork included AS events of ROBO1 and CLIP1, both of which had neural-regulated micro-exons (exons with 3□27 nt) involved in an AS interaction network associated with the autism spectrum disorder in the previous study [14]. Among our AS events, three micro-exon events (AP2M1, CLASP1, ROBO1) were detected as neural-regulated in the previous study. In particular, ROBO1 exon 18 skipping is known to induce helical domain exclusion and is involved in the loss-of-function of the ROBO1-SLIT2 signaling, thereby modulating neurogenesis and proliferation (Figure 4E). In our result, exon exclusion of ROBO1 was significant (permutation *P*-value = 0.001, dPSI = -0.265; Additional File 1: Table 1) and moderately correlated with the GSVA scores of the REACTOME ROBO receptor signaling pathway (*r* = -0.53). Meanwhile, exonic regions in our AS sets were involved in the protein domain (45.7 %) and isoform-specific interactions (21.3 %) (Additional File 2: Table S4). SE events by RBFOX1 knockdown induce an increase in the alteration of the protein domain, NMD, and repeat region, but decrease PTM and PPI.

### Performance comparison using SF3B1-associated MDS RNA-Seq dataset

The ability of the four different methods to detect DAS was evaluated by using the case study 1 database (details described in Methods). We selected an additional three programs, rMATS, MISO, and SUPPA2 for comparison [7–9]. We obtained four DAS sets from ASpediaFI (281 events at 194 genes), rMATS (596 events at 415 genes), MISO (685 events at 461 genes), and SUPPA2 (129 events and 99 genes) that were extracted from the results. To evaluate the functional enrichment of the detected DAS genes, we assessed the enrichment in the HM and expansion gene set, which are clinically known pathways regulated in MDS SF3B1 MUT samples [4,13,23–26]. The ASpediaFI result showed the best performance based on metrics like Fisher’s exact test *P*-value and *F*_*1*_ score (Figure 5A, Additional File 3: Figure S2).

**Figure 5.**
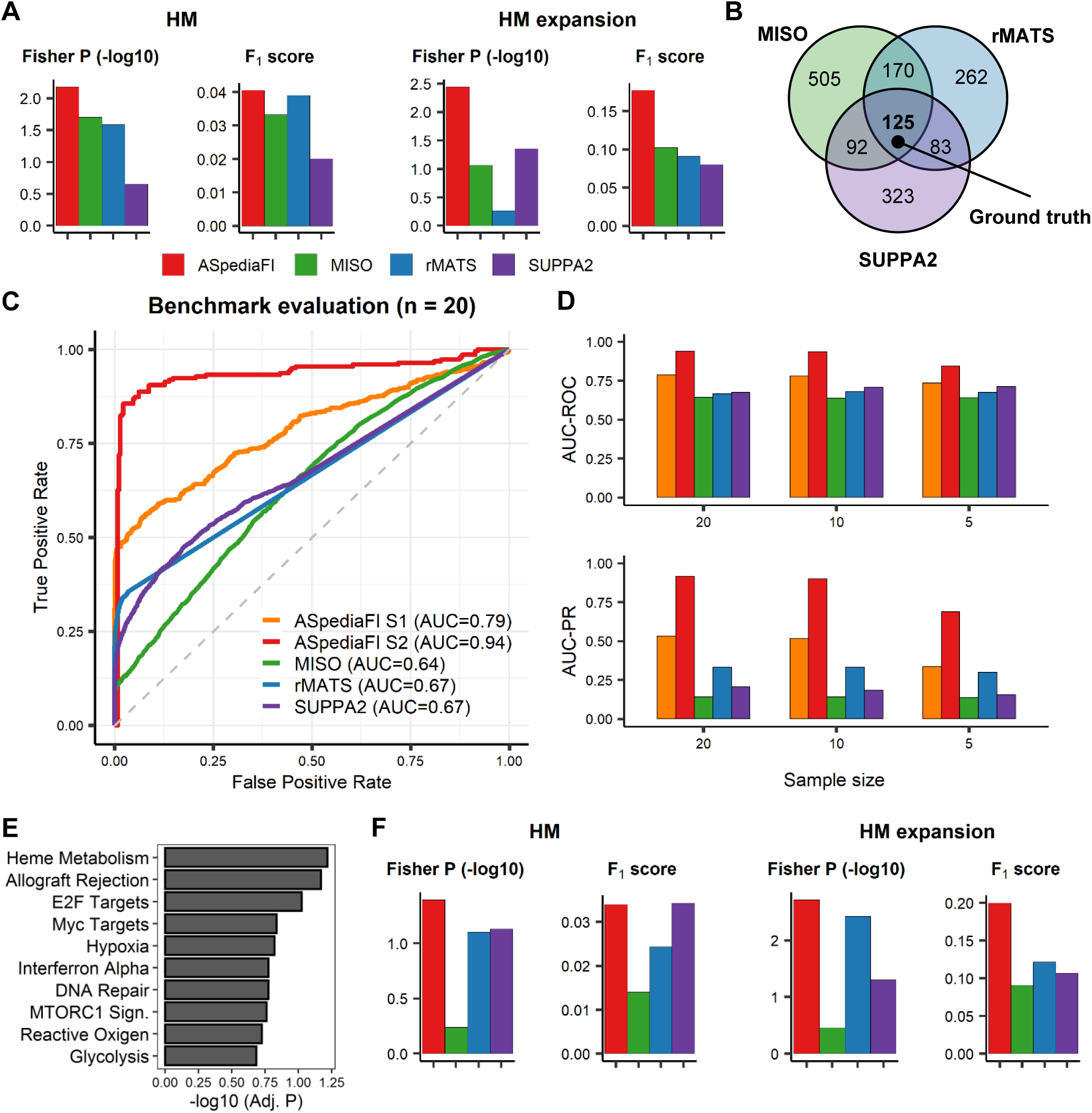
Performance evaluation of ASpediaFI and comparison with three other methods (MISO, rMATS, and SUPPA2) (A) Barplots of Fishers’ exact test *P*-values and *F*_*1*_ scores to test pathway enrichment for both HM and expansion sets. Enrichment was tested from two pathway gene sets, and AS event gene sets identified from four methods. (B) Venn diagram of DAS genes among three methods analyzing case study 1 MDS dataset. The intersecting DAS genes (n = 125) among all three methods serve as the ground truth for the simulated dataset. (C) ROC curves to evaluate the accuracy of four methods detecting DAS from a simulated dataset. ROC curves for each method illustrate true-positive rate (y-axis) against false-positive rate (x-axis). AUC values are described for each method. The dotted diagonal line corresponds to a ROC curve when DAS predictions are randomly guessed (AUC = 0.5). (D) Barplots of AUC-ROC and AUC-PR for the evaluation of sample size effect (n= 20, 10, 5 replicates per condition). Bar colors indicate the same method as in Figure 5C. (E) GSEA *P*-value barplot of highly ranked hallmark pathway from the simulated dataset that we imitate SF3B1 MUT and WT. HM pathway is detected on top. (F) Barplots of Fishers’ exact test *P*-values and *F*_*1*_ scores to test pathway enrichment for both HM and expansion sets. AS event sets were extracted from the simulated data analysis using four methods.

Between the two gene sets (Figure 5A), the ASpediaFI recall (0.175) in the HM expansion set was much better than that in HM (0.04). This result suggests that our method provides better performance for identifying AS events in a novel gene set like HM expansion compared to the other three tools (Figure 5A, Additional File 3: Figure S2). Meanwhile, to reduce the bias of comparing DAS sets with a different number of events, we modified the criteria for differential splicing such that the top 300 ranked AS events after filtering out events with | dPSI | < 0.1 are selected. This strategy substantially decreased the total counts in MISO and rMATS. ASpediaFI exhibited the best performance across Fisher’s exact test, precision, recall, and *F*_*1*_ than others (Additional File 2: Table S6). Regardless of the numbers of DAS events based on relaxed or strict thresholds, ASpediaFI consistently outperformed the other methods in detecting biologically relevant DAS events enriched in HM and expansion set.

### Performance evaluation on simulated datasets

To evaluate the ability of ASpediaFI to detect biologically relevant DAS events under a simulated environment, we generated a simulation dataset imitating the genomic characteristics of the MDS MUT and WT datasets. To artificially induce DAS events, we used the intersection DAS set identified by MISO (892 genes), rMATS (640 genes), and SUPPA2 (623 genes) as the ground truth for the evaluation (Figure 5B). Transcript counts of 20 replicates per condition were simulated from the distributions estimated by SF3B1 MUT and WT samples. We assigned the pre-determined relative isoform abundances for 125 ground truth AS genes collected from the intersection of three results, while those of other genes were drawn from the uniform distribution. ASpediaFI was excluded from generating these simulated RNA-Seq data for the blind test.

The ability of the four methods to detect previously defined ground truth AS events was verified. To measure the discriminative power, we computed AUC, AUC-ROC, and AUC-PR. As ASpediaFI runs DRaWR generating stationary probabilities for two stages, we used both stat-P’s for the comparison. The first stage (S1) stat-P values were computed for the whole AS events, and the final stat-P values (S2) were considered as refined ranks, enhancing the internal performance. ASpediaFI S1 achieved a higher AUC-ROC value of 0.79 than MISO (0.64), rMATS (0.67), and SUPPA2 (0.67) (Figure 5C). The performance difference manifested the overall false-positive rate (0□0.75). Not surprisingly, ASpediaFI S2 was better (AUC-ROC = 0.94) than S1. Moreover, we assessed the accuracy based on the lower number of samples per condition between the four methods. Compared to the fully simulated dataset (20 replicates per condition), the three other methods showed consistent performance for the smaller sample sizes (10 replicates) (Additional File 3: Figure S3). In the smallest dataset (n=5), our AUC-ROC decreased to 0.73 from 0.78, but the difference of true-positive rate was still maintained over the most important region (false-positive rate 0 ∼ 0.25). Although the AUC values of ASpediaFI slightly decreased, followed by sample size, S1 consistently exhibited superior performance compared to the other methods across the three simulated datasets of different sizes (Figure 5D, Additional File 3: S3). Additionally, the discriminative power of S2 remained reasonably stable under varying sample sizes.

We further examined the biological relevance of DAS events detected from the fully simulated dataset. As the simulation RNA-Seq samples were derived from SF3B1 MUT and WT samples, and as the ground truth AS events were defined based on the MDS sample analysis, we expected that the simulated samples would maintain the characteristics associated with the HM pathway dysregulation. Before DAS identification, we validated our assumption using GSEA with the gene expression profile. Finally, the HM pathway was consistently observed as the most significantly enriched pathway in the simulated dataset, as in case study 1 (adjusted *P*-value = 0.06; Figure 5E). The previously identified hypoxia and MTORC1 signaling pathways were also retained (adjusted *P*-value = 0.15, 0.17). Next, to make a fair comparison with ASpediaFI (DAS events n = 499), we identified the top 500 DAS after filtering | dPSI | > 0.1 using the other three methods. ASpediaFI exhibited a higher degree of enrichment in both the HM and expansion sets compared to the other three methods based on Fisher’s exact test *P*-value and *F*_*1*_ score (Figure 5F, Additional File 3: Figure S4). When stricter statistical cutoffs (FDR < 5% for rMATS and SUPPA2, Bayes Factor ≥ 5 for MISO) were applied to the other three methods, 210□280 AS events were identified (Additional File 2: Table S7). rMATS showed the highest enrichment according to Fisher’s exact test (HM *P*-value = 0.015, expansion *P*-value = 0.00085). Nevertheless, we observed that ASpediaFI showed better performance with respect to the *F*_*1*_ score (ASpediaFI 0.2, rMATS 0.107) of HM expansion than rMATS. It implies that our method detected novel AS events that are not present in the curated gene set. Overall, based on our benchmarking analyses using computationally simulated datasets, ASpediaFI showed a higher potential for identifying biologically-relevant DAS events.

## DISCUSSION

After the advent of next-generation sequencing, various novel methods for DAS analysis have been developed. Although approaches for DAS event identification have improved in accuracy, it is still a challenge to interpret the biological relevance as well as integration with regulatory mechanisms with DAS events. Here, we suggest an integrative method, ASpediaFI, to systematically identify AS events, co-expressed genes, and pathways regulated by the transcriptome. ASpediaFI ranks AS events, pathways, and genes, and also intuitively provides functional interactions in the form of an interaction network. It enables the users to understand global regulation and specific pathways by spliceosome and to choose more relevant AS events as markers.

In order to verify the intrinsic ability of ASpediaFI, we analyzed three case study datasets of MDS, STAD, and RBFOX1 knockdown. Pathway analysis results using our method presented remarkable consistency with GSEA or GSVA using the gene expression profile. This consistency can be attributed to the fact that our analysis starts with getting a query from the DEG set and performs RWR via a heterogeneous network that includes correlated AS with gene expression. Despite tumor heterogeneity in case 1, the high number of replication (total samples n = 84) facilitated the identification of AS events, their interacting genes, and pathway-level regulation by SF MUT. Next, we succeeded in identifying the gastric cancer EMT subtype based on the DAS. The subtype was revealed to be the one with the poorest survival among the four known gastric cancer subtypes [39]. Even though we identified a small size DAS set of around 200 events, our result demonstrated the discriminative power to classify samples by SF regulation (Figure 2C, Figure 3A). In particular, the three representative AS events, ENAH, FGFR2, and TCF7L2, that were identified only by ASpediaFI, had the potential to effectively classify the gastric cancer EMT subtype. It was comparable to the previous classification of the EMT subtype using the gene signature of over 300 genes [39]. In case 3, the previous RBFOX1 study performed GSEA and network-based module identification for each DEG and DAS sets [5]. This previous approach required the identification of relatively large DAS sets (n = 996). To uncover relevant biological process, the large size DAS set was divided into subsets by SE type or exon inclusion, and multiple sets were respectively used for GSEA. Moreover, independent analyses of DEG and DAS could not be interlinked to explain the systematic interactions between AS events, although the previous study successfully revealed the regulation of neuronal development by RBFOX1. Moreover, the pathway revealed using the two gene sets used in the previous study was complementary for uncovering neuronal development by RBFOX1, as already shown (Figure 4C). The multiple independent tests and complementary result highlight the advantage of our method.

In the case studies, our method correctly identified the AS type usage based on the role of SF with respect to recognizing donor and acceptor sites. In the previous study comparing several DAS methods, exon-based approaches mostly showed the best AU-ROC in terms of the SE event among the four AS types compared to the isoform-based methods [12]. Moreover, SE is the predominant type in the human gene model, and its PSI value calculated from three junctions and exons is more stable than A3 and A5 calculated from transcript regions narrower than SE. Therefore, exon-based DAS analysis applications have the potential to include bias according to AS type than isoform-based methods [12]. U2AF1, like SF3B1, is a member of the U2 complex and is known to recognize the 3′ dinucleotide motif AG, so A3 and RI could increase in the background of U2 complex member deficiency [6]. In case study 1, our result mirrors the characteristics of spliceosomes. Among the four tools we used, the AS type proportions of SUPPA2 resembled ours in case studies 1 and 2. The results of case study 2 and 3 were similar to previous results that identified the induction of SE events by ESRP1 and RBFOX1 [5,30,31]. In contrast to our result, rMATS most frequently detected SE events in the case studies. We deduced that the previous study on the MDS dataset had to carry out two comparisons with two different controls to avoid the SE bias [13]. When calculating the ratios of SE over the sum of A3 and A5 from several EMT-associated DAS results, rMATS (SE n=239, FDR < 10%; 18.8 times) and MADS+ (20 times) detected SE events at a higher frequency than previous analyses using Affymetrix exon 1.10 microarray (8.8 times) and RNA-Seq dataset considering sequence motif (3.6 times) and ours (3.1 times) [7,30,31,34]. That is, ASpediaFI provided results with a minimal bias toward SE, similar to SUPPA2.

To evaluate the performance of ASpediaFI and to compare it with other tools, we selected three analysis tools. In the early stage, we tried to add JUM, but we decided not to use it due to the extremely lower number of DAS passing the FDR threshold (< 5%). JUM can identify novel structured AS events not present in the transcriptome annotation [10]. We speculated that the advantage of JUM with respect to identifying novel events paradoxically reduced the detection of proper DAS events. Meanwhile, we chose the HM pathway as a gold standard for the performance evaluation of the analysis of the MDS dataset, based on the evidence from case study 1 and multiple previous clinical MDS studies [4,13,23–26]. The previous studies consistently reported the deficiency in heme biosynthesis and iron homeostasis due to splicing upon analyzing 12□100 samples. Unfortunately, the previous four splicing signatures identified from SF3B1 MUT samples did not have uniform quality, and identified AS signatures were small size (n = 20□202) except for one (n = 1403) [4,13,24,40]. However, we tried to perform Fisher’s exact test and Jaccard index for these four splicing signatures with our ASpediaFI AS results, Iron homeostasis transport, inflammatory response, HM and expansion set to evaluate the functional relevance based on previous studies [4,6,13,23–26]. ASpediaFI showed remarkable consistency (Fisher *P*-value < 0.0003) with three signatures except for the smallest sized signature (n = 20; *P*-value = 1). Next, the HM and expansion set represented the best enrichment (HM Fisher median *P*-value = 0.1; expansion *P*-value = 0.1) than others (Iron homeostasis transport *P*-value = 0.3; Inflammatory response *P*-value = 0.8). Based on these results and previous studies, we concluded that DAS events induced by SF3B1 MUT in MDS are enriched in the HM pathway and continued our evaluations.

During the evaluation using a simulated dataset, our method consistently showed the best performance compared to the other three tools. We tried to generate simulated RNA-Seq samples imitating actual MDS characteristics. We evaluated our capability to detect biologically relevant AS events in a dedicated setup, including the estimation of MUT and WT transcript count distributions and recurrent detection of DAS. Finally, we worked on benchmark evaluation as well as investigation of HM pathway enrichment. ASpediaFI generated the best ROC curves and presented a true-positive rate difference continuously across a long-range of false-positive rates (< 0.75) (Figure 5C). In the datasets with smaller sample sizes (n = 10 and 5), our method still showed the best result. For the evaluation of our tool, we used both S1 and S2 scores (Figure 5C). However, the second stage RWR is performed to rescore only AS events selected in S1. Therefore, S1, which is run on total AS events, is more suitable for comparison, and the outstanding achievement of S2 should be carefully interpreted. To achieve the best performance for each tool, we optimized parameters, such as FDR, dPSI, or BF and generally used cutoff values of other studies [7,10,24]. Sometimes, we removed additional filtering (dPSI) and only considered numerical scores (FDR or BF) from each tool. Despite these attempts, MISO demonstrated a weak performance in AUC-PR and HM enrichment. While rMATS showed the best performance in HM enrichment, ASpediaFI presented the best overall performance.

Our novel integrative approach using both PSI and gene expression offers a unique advantage. Instead of independent multiple GSEA tests for DAS and DEG, ASpediaFI systemically elucidates interactions between AS and genes and delineates pathway regulation. Another novel characteristic is its ability to identify relevant pathways using small size DAS sets. In contrast, other studies analyzed approximately 500□1000 AS events to reveal biological functions, and investigated pathways by dividing sets into inclusion and exclusion events. However, our method required fewer than 300 AS events to identify specific pathways in the three case studies. The total counts of our AS results are close to the recommended gene set size of at least 15 to at the most 200 genes [18] essential to identify splicing markers. Besides, there are additional advantages. AS event IDs of ASpediaFI results could be used to query the ASpedia database to explore comprehensive functional sequence features like protein domain, NMD, and isoform-specific interaction. Our tool has no dependency on any organisms or alignment tool. ASpediaFI refers dataset or file formats—BAM file, gene model, PPI, or gene sets—widely used in gene expression analyses. Moreover, our method supports fast execution time. The most time-consuming jobs to read bam files are provided with multi-thread option, and the principal analysis of DRaWR S1 and S2 except preprocessing is executable in a PC environment (RAM 16GB, CPU 3.40GHz and 2 minutes of execution time for case 1 SF3B1 dataset with total 82 RNA-Seq samples).

There are several limitations to ASpediaFI. Our method requires a reference interaction network and gene set. Prior to network establishment, our method involves filtering based on several criteria, including low gene expression and standard deviation of PSI. While it is effective at excluding unreliable PSI values calculated from lowly expressed genes, it is subject to the loss of lowly expressed true-positive AS events. As shown in the performance evaluation, our application needs at least five samples per condition to obtain a stable result. Additionally, ASpediaFI requires at least three samples per condition to calculate the correlation coefficient between the AS and gene. In a further development, we expect to improve the applicability of our method to a small dataset with less than five replicates or even without replication. Moreover, we also hope to extend our algorithm to the analysis of novel conditions like time-series or continuous statement of SF.

## CONCLUSION

In this study, we developed ASpediaFI and analyzed RNA-Seq datasets to verify the capability of our method to interpret biological processes regulated by splicing. As shown in the three case studies, ASpediaFI successfully identified AS events and relevant pathways involved in query DEGs. On comparison with three other three programs, ASpediaFI showed superior performance, as determined by the AUC-ROC and AUC-PR. We expect that ASpediaFI will uncover novel roles and global regulation of SFs.

## MATERIAL AND METHODS

### Data preparation

ASpediaFI requires input files, including a gene model, RNA-Seq BAM files, gene expression profiles, pathway gene sets, and a global gene-gene interaction network. First, AS events were identified using a gene model of a GTF file and classified into the following five types: alternative 5′ splice site (A5), alternative 3′ splice site (A3), skipping exon (SE), mutually exclusive exons (MXE), and retained intron (RI). PSI values of the identified events were calculated based on read counts mapped to exons and splice junctions. ASpediaFI uses these AS events, pathway gene sets, and a gene interaction network as reliable sources of interactions for the construction of a heterogeneous network. Our heterogeneous gene interaction network refers to a reference gene interaction. In our analysis, we collected and curated reference-based interaction databases (BIND, DIP, HPRD, and REACTOME) to build a reference human gene interaction compendium, which contains 10,647 genes and 54,037 interactions [19,41–43]. We also referred to public pathway databases (hallmark, REACTOME, and KEGG) and obtained a total of 910 human pathway gene sets [18,19,44].

### Heterogeneous network construction

Based on the biological information inferred from the RNA-Seq datasets and public databases, ASpediaFI constructed a heterogeneous network composed of gene nodes and two types of feature nodes: AS event and pathway. The heterogeneous network allows interactions between genes and between gene and feature node of gene-AS and gene-pathway. ASpediaFI refers to a reference network to connect gene interactions. Gene-gene interaction edges were weighted with the absolute value of the Pearson correlation coefficient calculated from gene expression. Gene-AS interaction edges are connected if the absolute value of the Spearman correlation coefficient between gene expression and PSI exceeds a user-defined threshold. Due to the nonlinear relationship between gene expression and PSI values, we used the Spearman correlation coefficient as a measure of association strength for gene-AS [45]. Finally, gene-pathway edges are weighted to 1 if the corresponding gene belongs to the corresponding pathway gene set.

### Query-specific subnetwork identification using DRaWR

To explore the important submodules, we employed DRaWR, which is the extension of random walk with restart (RWR) using a heterogeneous network consisting of feature nodes [20]. The DRaWR algorithm performs two-stage RWR in which a functional subnetwork is extracted in the first stage, and nodes in the subnetwork are ranked by associations with a query gene set in the second stage (Figure 1B).

Let *A* be an adjacency matrix representing our heterogeneous network. The adjacency matrix can be expressed as:

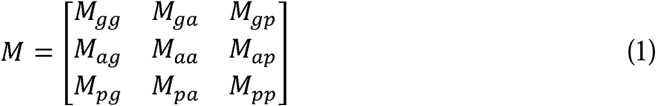

where submatrices *M*_*gg*_, *M*_*ga*_, and *M*_*gp*_ exhibit edges between gene-gene, gene-AS, and gene-pathway. Therefore, the entries of *M* can be written as:

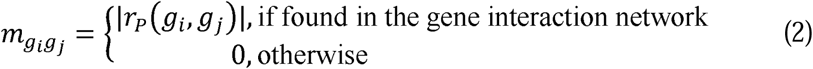

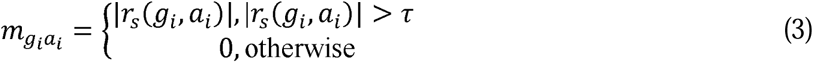

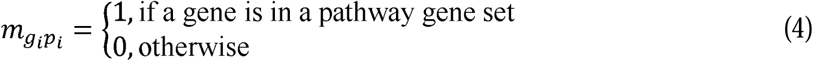

where *r*_*P*_ and *r*_*S*_ are the Pearson and Spearman correlation coefficients, respectively and *τ* is a user-defined threshold. Note that 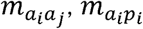, and 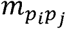 are all zero as there are no edges among feature nodes. Before running RWR, each nonzero submatrix is normalized such that its entries total 1, and the whole normalized adjacency matrix is again normalized by column to obtain a transition matrix *T*.

Given a transition matrix, the RWR algorithm can be formulated as:

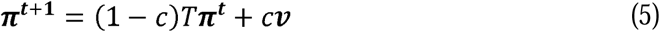

where 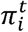 is the probability that the walker will stay at node *i* after the *t*th iteration, *c* is the probability of restart, and *v*_*j*_ is the probability of restarting at a node *j*. That is, for a query gene set *Q,v*_*j*_ is 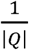 if *j*∈ *Q* and 0 otherwise. We assumed ***π***^**0**^ to be a uniform probability vector such that 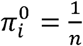, where *n* is the number of all nodes in a heterogeneous network.

In the first stage of DRaWR, RWR is run twice, once (Stage 1; S1) with a query gene set and another (Stage 2; S2) with all genes in the heterogeneous network as the restart set. The difference between the stationary probabilities (stat-P) in the two runs, say 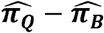, is a measure of relevance to a query gene set and used to rank AS event nodes and pathway nodes altogether.

Prior to stage 2, ASpediaFI extracts a query-specific subnetwork composed of gene nodes and the user-defined number of highly ranked AS event and pathway nodes. The adjacency matrix of the subnetwork can be expressed as:

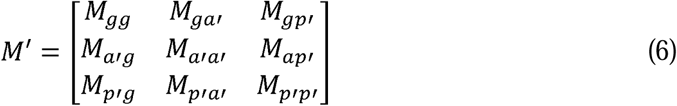

where *a*′ and *p*′ denote AS event and pathway nodes retained in the subnetwork. The second-stage RWR is performed on the subnetwork in the same way as stage 1 to calculate stat-P and produce final rankings of genes, pathways, and AS events.

### Evaluation of two-stage DRaWR and permutation test

ASpediaFI carries out a *k*-fold cross-validation at each stage of RWR to evaluate the performance of the DRaWR algorithm, in the same way as mentioned in the previous study [20]. A query gene set is partitioned into the user-defined number of subsets. For each subset, RWR is run with the remaining genes as the restart set to compute AUC (area under the curve) using the subset as true class labels and stat-P’s as predictions. In our analysis, we compared the average AUC at two stages for tuning parameters.

While the DRaWR algorithm removes feature nodes having low stat-P values under cutoff derived from querying a gene set before the second stage, all gene nodes are retained in the initial network and only provide their final relevance scores. In order to reduce the background effect of scoring and to filter out false positives, we included the permutation test on gene nodes in the evaluation procedure [46]. ASpediaFI runs *N* iterations of the second-stage random walks, in each of which a randomly sampled gene set of the same size as a query gene set is used as the restart set. The permutation *P*-value of gene node *i* is

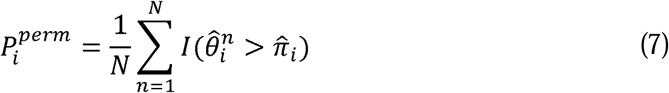

where 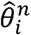 is the second-stage stat-P of node *i* when a randomly sampled gene set is given as a query, and *I* is an indicator function which gives 1 if 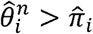 and 0 otherwise. ASpediaFI refers to stat-Ps as a score for ranking and selecting feature nodes, and permutation *P*-values for choosing pathway-related genes.

### RNA-Seq dataset preparation for case studies

#### Case study 1

The first case study was an RNA-Seq dataset (GEO accession number: GSE114922) from bone marrow-derived CD34+ hematopoietic progenitor cells of 84 patients with myelodysplastic syndrome (MDS) [13]. Patients exhibited hotspot mutations in three SF SF3B1 (n = 28), SRSF2 (n = 6), and U2AF1 (n = 8). We first assessed the quality of reads using FastQC v0.11.5, and aligned to the GRCh38 genome and the reference gene model GENCODE v31 using STAR v2.6.1b to follow the GDC pipeline with customized options: outFilterType = BySJout, alignEndsType = EndToEnd, alignSoftClipAtReferenceEnds = No, alignIntronMax = 10000, alignMatesGapMax = 10000) [47]. Gene expression profile was evaluated by RSEM v1.3.0 [48]. We calculated the PSI (percent spliced-in) profile from BAM files based on AS events derived from the input gene model. To extract the query gene set, differential expression analysis between the mutated and wild-type samples was performed using limma v3.42.0 [49]. The ASpediaFI analysis was run with the following options:

- restart (restart probability): 0.7
- num.folds (number of folds for cross-validation): 5
- num.feats (number of features to be retained in a subnetwork): 300
- low.expr (threshold average FPKM of genes): 1
- low.var (threshold variance of AS events): NULL
- prop.na (threshold proportion of missing PSI values): 0.05
- prop.extreme (threshold proportion of extreme PSI values – 0 or 1): 1
- cor.threshold (threshold Spearman’s correlation coefficient between genes and AS events): 0.4.

Based on this, we reconstructed three AS-gene interaction subnetworks regulated by three SF mutations from the second stage result of DRaWR. Additionally, highly-scored genes (permutation *P*-values < 0.05) were selected along with neighboring AS event nodes.

We further investigated the characteristics and biological relevance of the identified AS events. First, we classified the MDS samples into two groups, SF WT and MUT using the PSI profiles of identified DAS events. We performed hierarchical clustering with complete linkage on the Euclidean distance matrix of the PSI profiles to evaluate the discriminative performance, confirmed by principal component analysis (PCA). Next, we used rMATS to detect DAS between SF3B1 MUT and WT and compared it with our result. Based on previous MDS study analysis condition, we setup rMATS cutoffs (Cond1: | dPSI | > 0.1 & FDR < 0.05) [13]. The number of DAS identified by rMATS Cond1 is over twice of our AS result. To make similar condition, we additionally performed rMATS of more stringent cutoff conditions. We first applied the same thresholds (Cond1: | dPSI | > 0.1 & FDR < 0.05) to follow the methodology used in the previous study. The second thresholds (Cond2: | dPSI | > 0.1 & FDR < 0.0001) were determined so that the number of DAS events was similar to the ASpediaFI result. As our method refers to PPI genes to identify all interactions, only AS events in genes in our PPI compendium were considered to reduce the bias introduced using different background genes. We used the Fisher’s exact test and Jaccard index to measure how the results of ASpediaFI, rMATS Cond1, and Cond2 are enriched in the heme metabolism (HM) pathway, which was highly ranked in the previous study. Additionally, we defined a novel HM gene set ‘HM expansion set’ to test whether AS events interact with genes in the HM pathway and participate in the corresponding biological process. The HM expansion set included both HM genes and their neighbor genes derived from our gene interaction network. Fisher’s exact test and Jaccard index were also computed for the HM expansion set. To investigate the functional importance of AS genomic regions, we interrogated protein domain, NMD, and other sequential features of AS events using the ASpedia database for ASpediaFI, rMATS, Cond1, and Cond2 [22].

#### Case study 2

We chose the TCGA stomach adenocarcinoma (STAD) level 3 RNA-Seq dataset as another real dataset to investigate AS events and biological processes associated with ESRP1, a key splicing factor that regulates epithelial-mesenchymal transition (EMT) across multiple cancer types [2,7,50]. Of the 415 STAD patients, the highest and lowest 10% mRNA expression samples of ESRP1 were classified as ESRP1-high and ESRP1-low groups, respectively. Due to the absence of BAM files, we used SUPPA2 v2.3, as was done in the previous study, and we also used a gene model referred UCSC known genes to generate PSI profiles [51]. Statistical test for differential expression between the two groups was performed using limma to obtain a query gene set. We conducted the ASpediaFI analysis with the following options: restart = 0.7, num.folds = 5, num.feats = 300, low.expr = 1, low.var = NULL, prop.na = 0.05, prop.extreme = 1 and cor.threshold = 0.5. To compare our result, we performed DAS analysis using SUPPA2 diffSplice with the following options: nan-threshold = 10, area = 1000 and lower-bound = 0.05 [8]. SUPPA2 DAS set was obtained by selecting AS events with | dPSI | > 0.1 and adjusted *P*-value < 0.1. Next, we extracted an EMT-associated subnetwork from the final stage produced by DRaWR. To decrease the network size, we filtered out gene nodes with permutation *P*-values not less than 0.05.

Similarly, we tested the discriminative power of our DAS events by classifying STAD samples based on the Euclidean distance matrix of their PSI profiles using hierarchical clustering with average linkage. Meanwhile, we test how much our pathway result identified by ASpediaFI is consistent with GSEA analysis using gene expression profile. Our pathway result was collected by rankings determined by ASpediaFI. For analysis result using gene expression, we calculated sample-level pathway activity scores executing gene set variation analysis (GSVA) [32]. Difference of GSVA scores between high and low groups was tested by Wilcoxon rank-sum test. Next, we compared ASpediaFI with other DAS test method. The results from the two applications, ASpediaFI and SUPPA2, were compared using Venn diagram, Fishers’ exact test, and Jaccard index calculated from five EMT or ESRP1-associated splicing gene signatures [31,33–35]. Like in case study 1, AS event sets for two conditions were chosen to overlap with global PPI genes. As in the first case study, we compared the sequential features of AS events detected by ASpediaFI and SUPPA2 by retrieving from the ASpedia database.

#### Case study 3

The last RNA-Seq data (GEO accession number: GSE36710) comprised five replicates of the shRBFOX1 (RBFOX1 knockdown) and shGFP (control) cell lines [5]. Single-end RNA-Seq reads aligned to the GRCh37 genome and the reference gene model Ensembl v71 using STAR v2.6.1b with the same options as in case study 1. We calculated gene expression and PSI values using RSEM and our quantification tool. A query gene set was obtained from the DEG test between RBFOX1 knockdown and control groups using limma. The following options were selected for the ASpediaFI workflow: restart = 0.7, num.folds = 5, num.feats = 300, low.expr = 1, low.var = NULL, prop.na = 0.05, prop.extreme = 1, cor.threshold = 0.8. We then interrogated an RBFOX1-regulated subnetwork. From the final network produced by the DRaWR algorithm, we retained gene nodes with permutation *P*-values less than 0.05 and their neighboring AS event nodes.

To examine the enrichment of our AS genes in known neuronal genes, we calculated the Jaccard index between our AS gene set and known gene signatures, as done in the previous study [5]. We prepared three published gene signatures containing genes inferred from transcriptomic analysis of RBFOX1 and RBFOX2, and those showing RBFOX1-dependent splicing in autism spectrum disorder (ASD) brains [52–54]. We also compared with three control gene signatures – mitochondria, epilepsy, and ataxia [5]. To evaluate the performance of ASpediaFI for identifying biologically relevant pathways, we performed gene set enrichment analysis (GSEA) on gene nodes with permutation *P*-values less than 0.05 using DAVID v6.8 [55]. Our GSEA result was compared with previously identified two gene sets: blue module and DAS [5]. The blue module comprising 737 genes is a subnetwork identified by WGCNA using gene expression profiles; DAS contained 603 differentially spliced genes detected by DESeq [56]. We also explored the sequential features of our AS events retrieved from the ASpedia database.

### Performance comparison using SF3B1 mutation MDS patients RNA-Seq dataset

To compare the performance of ASpediaFI against other widely used DAS detection tools, we extended case study 1 using the SF3B1-associated MDS dataset. In addition to rMATS v4.0.2, we applied MISO v0.5.4 and SUPPA2 v2.3 to the same MDS RNA-Seq dataset [7–9,13]. We customized settings for DAS analysis to reflect the characteristics of each tool. rMATS analysis results were collected from case study 1 and additional cutoffs (| dPSI | > 0.1 and FDR < 5%) were applied. For MISO, as only pairwise comparisons are allowed in DAS analysis, we merged BAM files for multiple samples per condition (SF3B1 MUT: 28 cases and WT: 56 controls). DAS analysis was performed using the pooled version of BAM files, and other parameters were used in default settings. We, therefore, filtered the resultant DAS events with more stringent minimal coverage and Bayes factor (BF) than default values (BF ≥ 20, the sum of inclusion and exclusion reads ≥ 300, at least 30 inclusion and exclusion reads), and eliminated AS events by the same cutoff (| dPSI | ≤ 0.1) with rMATS. For SUPPA2, to obtain PSI profiles, we quantified transcript expression in TPM units using RSEM v1.3.0. Next, we executed the embedded modules *psiPerEvent* to generate PSI profiles and *diffSplice* to detect DAS events using default options. The same thresholds with rMATS were also applied to select final DAS events derived from SUPPA2. To evaluate the performance of the four methods, we tested gene set enrichment of HM pathway referring to the top-ranking results in case study 1 and previously published studies [4,13,23–26]. To test the enrichment of DAS events for each tool, we converted DAS events to gene symbols and computed Fisher’s exact test *P*-values and F1 scores for HM and expansion pathway gene sets.

### Performance benchmark using simulated datasets

To evaluate the ability to detect functionally-enriched DAS of ASpediaFI, we generated a simulation dataset close to the actual MDS patient RNA-Seq dataset. To define the ground truth AS gene set for simulation, we intended to select AS genes that are highly likely to occur for the real MDS samples instead of randomly chosen genes. Therefore we investigated gene sets using three different methods. We applied the same running options for rMATS, MISO, and SUPPA2 as with previous evaluations using MDS samples. To identify more DAS genes on intersection set, we imposed relatively less stringent cutoffs: | dPSI | > 0.025, FDR < 10% for rMATS and SUPPA2. For MISO, additional strict thresholds were applied to balance the number of DAS events with the other two methods (BF ≥ 800, the sum of inclusion and exclusion reads ≥ 700), as well as the same relaxed cutoff (| dPSI | > 0.025). Finally, the resulting set of DAS genes overlapping between the three tools was assigned as our ground truth for the simulated dataset.

Next, we generated 20 replicates per condition (SF3B1 MUT and WT) via Flux Simulator [57], executing scripts from the previous simulation study [12]. To simulate realistic RNA-Seq reads, we referred to the real RNA-Seq samples of MDS patients with SF3B1 MUT and WT. Transcript counts were sampled from a negative binomial distribution with mean and variance estimated for MUT and WT conditions of the original MDS BAM files. For the deliberately chosen true DAS genes, we set relative isoform abundances such that the last isoform took a pre-determined proportion (0.8 for MUT and 0.2 for WT), while others equally shared the rest. Isoform-level abundances of other genes were drawn from a uniform distribution. The simulated RNA-Seq reads for each replicate with mean base coverage of 65 were then mapped to the GRCh38 genome along with the GENCODE v31 gene model, using STAR v2.5.1b. Additionally, to evaluate the effect of sample size, 10 and 5 replicates per condition were randomly chosen from the full simulated dataset. We performed DAS tests using the simulation dataset for ASpediaFI and three other methods. ASpediaFI was run with the following options: restart = 0.7, num.folds = 5, num.feats = 500, low.expr = 1, low.var = NULL, prop.na = 0.05, prop.extreme = 1. For each simulated datasets of different sizes (n=20, 10, 5 replicates per condition), the cor.threshold option was adjusted by the number of detected AS event nodes (0.4, 0.5, 0.8, respectively). The three tools were applied using options previously described. As the genome-wide ranking results were compared, additional filtering by dPSI was excluded.

To evaluate the accuracy of the four methods, we generated receiver operating characteristic (ROC) curve and computed the area under the curve (AUC-ROC) metric, using R PRROC package [58]. We also calculated the area under the precision-recall curve (AUC-PR) metric. Ranking of AS events were computed based on measures of 1 – adjusted *P* values for rMATS and SUPPA2, BF for MISO, and stat-P for ASpediaFI S1 and S2 were provided. Moreover, to assess the effect of sample size, we computed AUC-ROC and AUC-PR metrics using the simulated datasets of randomly-chosen smaller sample sizes (n = 10, 5 replicates per condition).

For further performance evaluation, we investigated pathway enrichment, and evaluated whether the four methods maintained their ability to identify biologically-relevant AS events using the simulated dataset. Based on previous studies and case study 1, we assumed that the HM pathway is dysregulated in MDS patients with SF3B1 mutation [4,13,23–26]. As our simulation dataset was derived from the actual MDS patient sample analysis result, we investigated the pathway status similar to the previously described process. To confirm GSEA consistency between DAS and DEG, we applied GAGE to perform GSEA using gene expression profiles for hallmark pathways [59]. Next, for the DAS enrichment test, we extracted an equal number (top 500) of most significant DAS events for each tool after filtering out by |dPSI| ≤ 0.1. Finally, we assessed the enrichment of AS event sets for the four methods by conducting Fisher’s exact test and computing the *F*_*1*_ score from HM and expansion sets.

## Abbreviations

AS: Alternative splicing
DRaWR: Discriminative random walk with restart
DAS: Differential alternative splicing
DEG: Differentially expressed genes
EMT: Epithelial-to-mesenchymal
GSEA: Gene set enrichment analysis
PSI: Percent spliced in
SF: Splicing factor
Stat-P: Stationary probability

## Declarations

### Ethics approval and consent to participate

Ethics approval was not applicable for this study.

### Competing interests

The authors declare that they have no competing interests.

### Availability of data and materials

Datasets used in this manuscript are accessible at GEO (accession: GSE114922 and GSE36710) and GDC (TCGA STAD RNA-Seq level3). ASpediaFI is supported as an R package open source program. The tool, user manual and case study are publicly available at Bioconductor (https://bioconductor.org/packages/ASpediaFI).

### Authors’ contribution

Conceptualization, supervision and funding acquisition, C.P.; algorithm implementation and analysis, D.Y., K.L., and D.H.; evaluation, D.Y. and K.L; writing, review, and editing, D.Y., K.L. S.Y.C., and C.P.

### Funding

This work was supported by National Research Foundation of Korea grant funded by the Korea government (NRF-2019R1A2C1003401); National Cancer Center Grant (NCC-1910040).

## Notes

### Competing Interest Statement

The authors have declared no competing interest.

## REFERENCES

1. Yang X, Coulombe-Huntington J, Kang S, Sheynkman GM, Hao T, Richardson A, et al. Widespread Expansion of Protein Interaction Capabilities by Alternative Splicing. Cell. 2016;164:805–17.

2. Saha A, Kim Y, Gewirtz ADH, Jo B, Gao C, McDowell IC, et al. Co-expression networks reveal the tissue-specific regulation of transcription and splicing. Genome Res. 2017;27:1843–58.

3. Salomonis N, Schlieve CR, Pereira L, Wahlquist C, Colas A, Zambon AC, et al. Alternative splicing regulates mouse embryonic stem cell pluripotency and differentiation. Proc Natl Acad Sci U S A. 2010;107:10514–9.

4. Dolatshad H, Pellagatti A, Liberante FG, Llorian M, Repapi E, Steeples V, et al. Cryptic splicing events in the iron transporter ABCB7 and other key target genes in SF3B1-mutant myelodysplastic syndromes. Leukemia. 2016;30:2322–31.

5. Fogel BL, Wexler E, Wahnich A, Friedrich T, Vijayendran C, Gao F, et al. RBFOX1 regulates both splicing and transcriptional networks in human neuronal development. Hum Mol Genet. 2012;21:4171–86.

6. Seiler M, Peng S, Agrawal AA, Palacino J, Teng T, Zhu P, et al. Somatic Mutational Landscape of Splicing Factor Genes and Their Functional Consequences across 33 Cancer Types. Cell Rep. 2018;23:282-296.e4.

7. Shen S, Park JW, Lu Z, Lin L, Henry MD, Wu YN, et al. rMATS: Robust and flexible detection of differential alternative splicing from replicate RNA-Seq data. Proc Natl Acad Sci. 2014;111:E5593–601.

8. Trincado JL, Entizne JC, Hysenaj G, Singh B, Skalic M, Elliott DJ, et al. SUPPA2: fast, accurate, and uncertainty-aware differential splicing analysis across multiple conditions. Genome Biol. 2018;19:40.

9. Katz Y, Wang ET, Airoldi EM, Burge CB. Analysis and design of RNA sequencing experiments for identifying isoform regulation. Nat Methods. 2010;7:1009–15.

10. Wang Q, Rio DC. JUM is a computational method for comprehensive annotation-free analysis of alternative pre-mRNA splicing patterns. Proc Natl Acad Sci. 2018;115:E8181–90.

11. Saraiva-Agostinho N, Barbosa-Morais NL. psichomics: graphical application for alternative splicing quantification and analysis. Nucleic Acids Res. 2019;47:e7–e7.

12. Liu R, Loraine AE, Dickerson JA. Comparisons of computational methods for differential alternative splicing detection using RNA-seq in plant systems. BMC Bioinformatics. 2014;15:364.

13. Pellagatti A, Armstrong RN, Steeples V, Sharma E, Repapi E, Singh S, et al. Impact of spliceosome mutations on RNA splicing in myelodysplasia: dysregulated genes/pathways and clinical associations. Blood. 2018;132:1225–40.

14. Irimia M, Weatheritt RJ, Ellis JD, Parikshak NN, Gonatopoulos-Pournatzis T, Babor M, et al. A highly conserved program of neuronal microexons is misregulated in autistic brains. Cell. 2014;159:1511–23.

15. Lee SCW, Abdel-Wahab O. Therapeutic targeting of splicing in cancer. Nat. Med. 2016. p. 976–86.

16. Warzecha CC, Sato TK, Nabet B, Hogenesch JB, Carstens RP. ESRP1 and ESRP2 Are Epithelial Cell-Type-Specific Regulators of FGFR2 Splicing. Mol Cell. 2009;33:591–601.

17. Wang B-D, Ceniccola K, Hwang S, Andrawis R, Horvath A, Freedman JA, et al. Alternative splicing promotes tumour aggressiveness and drug resistance in African American prostate cancer. Nat Commun. 2017;8:15921.

18. Liberzon A, Birger C, Thorvaldsdóttir H, Ghandi M, Mesirov JP, Tamayo P. The Molecular Signatures Database Hallmark Gene Set Collection. Cell Syst. 2015;1:417–25.

19. Croft D, O’Kelly G, Wu G, Haw R, Gillespie M, Matthews L, et al. Reactome: a database of reactions, pathways and biological processes. Nucleic Acids Res. 2011;39:D691–7.

20. Blatti C, Sinha S. Characterizing gene sets using discriminative random walks with restart on heterogeneous biological networks. Bioinformatics. 2016;32:2167–75.

21. Valdeolivas A, Tichit L, Navarro C, Perrin S, Odelin G, Levy N, et al. Random walk with restart on multiplex and heterogeneous biological networks. Bioinformatics. 2019;35:497–505.

22. Hyung D, Kim J, Cho SY, Park C. ASpedia□: a comprehensive encyclopedia of human alternative splicing. Nucleic Acids Res. 2018;46:58–63.

23. Pellagatti A, Cazzola M, Giagounidis AAN, Malcovati L, Della Porta MG, Killick S, et al. Gene expression profiles of CD34+ cells in myelodysplastic syndromes: Involvement of interferon-stimulated genes and correlation to FAB subtype and karyotype. Blood. 2006;108:337–45.

24. Dolatshad H, Pellagatti A, Fernandez-Mercado M, Yip BH, Malcovati L, Attwood M, et al. Disruption of SF3B1 results in deregulated expression and splicing of key genes and pathways in myelodysplastic syndrome hematopoietic stem and progenitor cells. Leukemia. 2015;29:1092–103.

25. Conte S, Katayama S, Vesterlund L, Karimi M, Dimitriou M, Jansson M, et al. Aberrant splicing of genes involved in haemoglobin synthesis and impaired terminal erythroid maturation in SF3B1 mutated refractory anaemia with ring sideroblasts. Br J Haematol. 2015;171:478–90.

26. Shiozawa Y, Malcovati L, Gallì A, Sato-Otsubo A, Kataoka K, Sato Y, et al. Aberrant splicing and defective mRNA production induced by somatic spliceosome mutations in myelodysplasia. Nat Commun. 2018;9:3649.

27. de Oliveira RM, Vicente Miranda H, Francelle L, Pinho R, Szegö ÉM, Martinho R, et al. The mechanism of sirtuin 2–mediated exacerbation of alpha-synuclein toxicity in models of Parkinson disease. PLoS Biol. 2017;15.

28. Nakamura K, Kageyama S, Yue S, Huang J, Fujii T, Ke B, et al. Heme oxygenase-1 regulates sirtuin-1–autophagy pathway in liver transplantation: From mouse to human. Am J Transplant. 2018;18:1110–21.

29. Shao J, Grammatikakis N, Scroggins BT, Uma S, Huang W, Chen JJ, et al. Hsp90 regulates p50cdc37 function during the biogensis of the active conformation of the hemeregulated eIF2α kinase. J Biol Chem. 2001;276:206–14.

30. Warzecha CC, Shen S, Xing Y, Carstens RP, Warzecha CC, Shen S, et al. The epithelial splicing factors ESRP1 and ESRP2 positively and negatively regulate diverse types of alternative splicing events Claude. RNA Biol. 2009;6:546–62.

31. Shapiro IM, Cheng AW, Flytzanis NC, Balsamo M, Condeelis JS, Oktay MH, et al. An emt-driven alternative splicing program occurs in human breast cancer and modulates cellular phenotype. PLoS Genet. 2011;7:e1002218.

32. Hänzelmann S, Castelo R, Guinney J. Open Access GSVA□: gene set variation analysis for microarray and RNA-Seq data. BMC Bioinformatics. 2013;14.

33. Yang Y, Park JW, Bebee TW, Warzecha CC, Guo Y, Shang X, et al. Determination of a Comprehensive Alternative Splicing Regulatory Network and Combinatorial Regulation by Key Factors during the Epithelial-to-Mesenchymal Transition. Mol Cell Biol. 2016;36:1704–19.

34. Warzecha CC, Jiang P, Amirikian K, Dittmar KA, Lu H, Shen S, et al. An ESRP-regulated splicing programme is abrogated during the epithelial–mesenchymal transition. EMBO J. 2010;29:3286–300.

35. Dittmar KA, Jiang P, Park JW, Amirikian K, Wan J, Shen S, et al. Genome-wide determination of a broad ESRP-regulated posttranscriptional network by high-throughput sequencing. Mol Cell Biol. 2012;32:1468–82.

36. Wieczorek K, Wiktorska M, Sacewicz-Hofman I, Boncela J, Lewiński A, Kowalska MA, et al. Filamin A upregulation correlates with Snail-induced epithelial to mesenchymal transition (EMT) and cell adhesion but its inhibition increases the migration of colon adenocarcinoma HT29 cells. Exp Cell Res. 2017;359:163–70.

37. Di Modugno F, Iapicca P, Boudreau A, Mottolese M, Terrenato I, Perracchio L, et al. Splicing program of human MENA produces a previously undescribed isoform associated with invasive, mesenchymal-like breast tumors. Proc Natl Acad Sci U S A. 2012;109:19280–5.

38. Weise A, Bruser K, Elfert S, Wallmen B, Wittel Y, Wöhrle S, et al. Alternative splicing of Tcf7l2 transcripts generates protein variants with differential promoter-binding and transcriptional activation properties at Wnt/beta-catenin targets. Nucleic Acids Res. 2010;38:1964–81.

39. Cristescu R, Lee J, Nebozhyn M, Kim KM, Ting JC, Wong SS, et al. Molecular analysis of gastric cancer identifies subtypes associated with distinct clinical outcomes. Nat Med. 2015;21:449–56.

40. Papaemmanuil E, Cazzola M, Boultwood J, Malcovati L, Vyas P, Bowen D, et al. Somatic SF3B1 mutation in myelodysplasia with ring sideroblasts. N Engl J Med. 2011;365:1384–95.

41. Bader GD, Betel D, Hogue CW V. BIND: the Biomolecular Interaction Network Database. Nucleic Acids Res. 2003;31:248–50.

42. Xenarios I, Rice DW, Salwinski L, Baron MK, Marcotte EM, Eisenberg D. DIP: the database of interacting proteins. Nucleic Acids Res. 2000;28:289–91.

43. Keshava Prasad TS, Goel R, Kandasamy K, Keerthikumar S, Kumar S, Mathivanan S, et al. Human Protein Reference Database--2009 update. Nucleic Acids Res. 2009;37:D767–72.

44. Kanehisa M, Sato Y, Kawashima M, Furumichi M, Tanabe M. KEGG as a reference resource for gene and protein annotation. Nucleic Acids Res. 2016;44:D457–62.

45. de Winter JCF, Gosling SD, Potter J. Comparing the pearson and spearman correlation coefficients across distributions and sample sizes: A tutorial using simulations and empirical data. Psychol Methods. 2016;21:273–90.

46. Zhu L, Su F, Xu YC, Zou Q. BBA - Molecular Basis of Disease Network-based method for mining novel HPV infection related genes using random walk with restart algorithm. BBA - Mol Basis Dis. 2018;1864:2376–83.

47. Dobin A, Davis CA, Schlesinger F, Drenkow J, Zaleski C, Jha S, et al. STAR: ultrafast universal RNA-seq aligner. Bioinformatics. 2013;29:15–21.

48. Li B, Dewey CN. RSEM: accurate transcript quantification from RNA-Seq data with or without a reference genome. BMC Bioinformatics. 2011;12:323.

49. Ritchie ME, Phipson B, Wu D, Hu Y, Law CW, Shi W, et al. limma powers differential expression analyses for RNA-sequencing and microarray studies. 2015;43.

50. Network TCGAR. Comprehensive molecular characterization of gastric adenocarcinoma. Nature. 2014;513:202–9.

51. Sebestyén E, Singh B, Miñana B, Pagès A, Mateo F, Pujana MA, et al. Large-scale analysis of genome and transcriptome alterations in multiple tumors unveils novel cancer-relevant splicing networks. Genome Res. 2016;26:732–44.

52. Zhang C, Zhang Z, Castle J, Sun S, Johnson J, Krainer AR, et al. Defining the regulatory network of the tissue-specific splicing factors Fox-1 and Fox-2. Genes Dev. 2008;22:2550–63.

53. Yeo GW, Coufal NG, Liang TY, Peng GE, Fu XD, Gage FH. An RNA code for the FOX2 splicing regulator revealed by mapping RNA-protein interactions in stem cells. Nat. Struct. Mol. Biol. 2009. p. 130–7.

54. Voineagu I, Wang X, Johnston P, Lowe JK, Tian Y, Horvath S, et al. Transcriptomic analysis of autistic brain reveals convergent molecular pathology. Nature. 2011;474:380–6.

55. Huang DW, Sherman BT, Lempicki RA. Systematic and integrative analysis of large gene lists using DAVID bioinformatics resources. Nat Protoc. 2009;4:44–57.

56. Anders S, Huber W. Differential expression analysis for sequence count data. Genome Biol. 2010;11.

57. Griebel T, Zacher B, Ribeca P, Raineri E, Lacroix V, Guigó R, et al. Modelling and simulating generic RNA-Seq experiments with the flux simulator. Nucleic Acids Res. 2012;40:10073–83.

58. Grau J, Grosse I, Keilwagen J. PRROC: computing and visualizing precision-recall and receiver operating characteristic curves in R. Bioinformatics. 2015;31:2595–7.

59. Luo W, Friedman MS, Shedden K, Hankenson KD, Woolf PJ. GAGE: Generally applicable gene set enrichment for pathway analysis. BMC Bioinformatics. 2009;10:161.

