## Supplementary material for "ASpediaFI: Functional interaction analysis of alternative splicing events": Figure S1-S4

**Figure S1 Exonic structures and protein domains of three EMT-associated AS events**

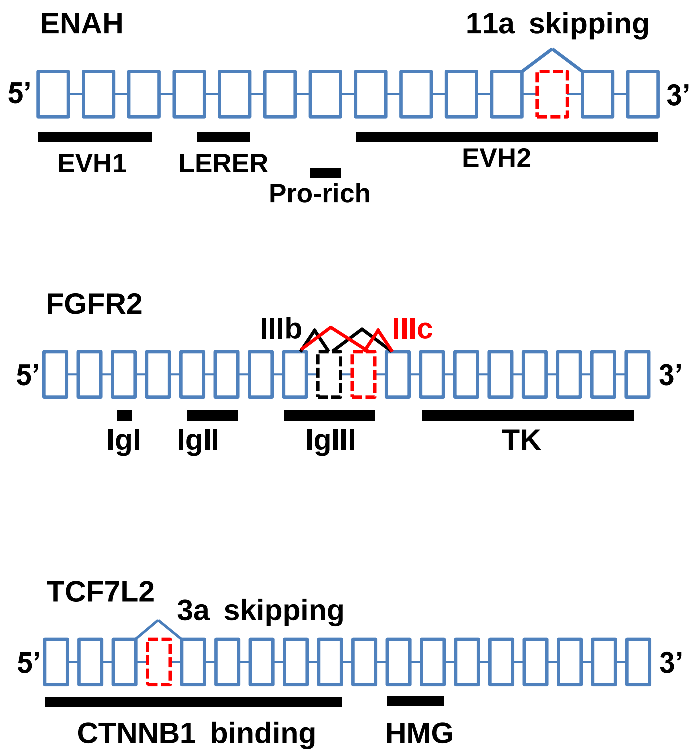

**Figure S2. Precision and recall for HM and HM expansion gene set calculated from SF3B1-MDS dataset analysis using four DAS detection tools.** The same resulting DAS genes from the comparison analysis (Figure 5A) were evaluated here. For HM pathway, ASpediaFI showed the highest value of precision among four tools, at the cost of relatively lower value of recall. For HM expansion gene set, ASpediaFI achieved higher precision and recall values compared to the other three methods. Each bar on the x-axis represents a DAS detection tool with its corresponding color at the legend (Bottom).

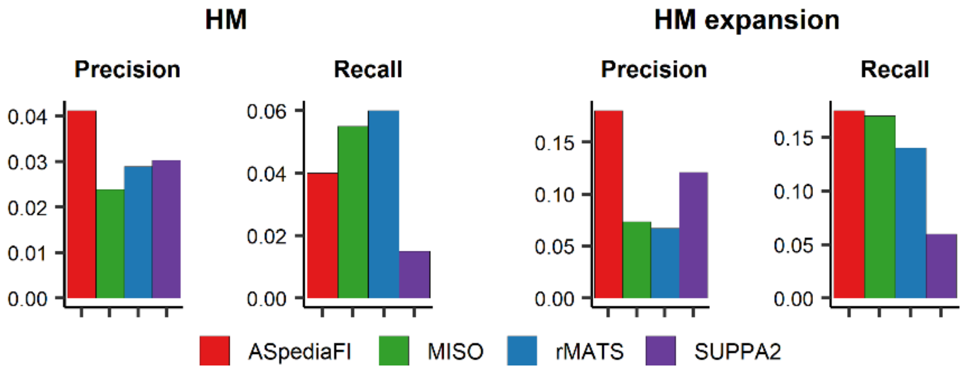

**Figure S3. ROC curves of four DAS analysis tools using simulated datasets of size 10 and 5 replicates per condition.** Benchmarking analyses were performed to measure the sample size effect. *n* replicates per condition (*n*=10, 5) were randomly subsampled from the original simulated dataset (*n*=20). Both stages of ASpediaFI S1 and S2 were evaluated. For each tool, the value of AUC metric and its corresponding color are displayed in the legend (bottom right). The dashed line indicates a non-discriminative method that randomly guesses true DAS events, and the corresponding AUC value is 0.5.

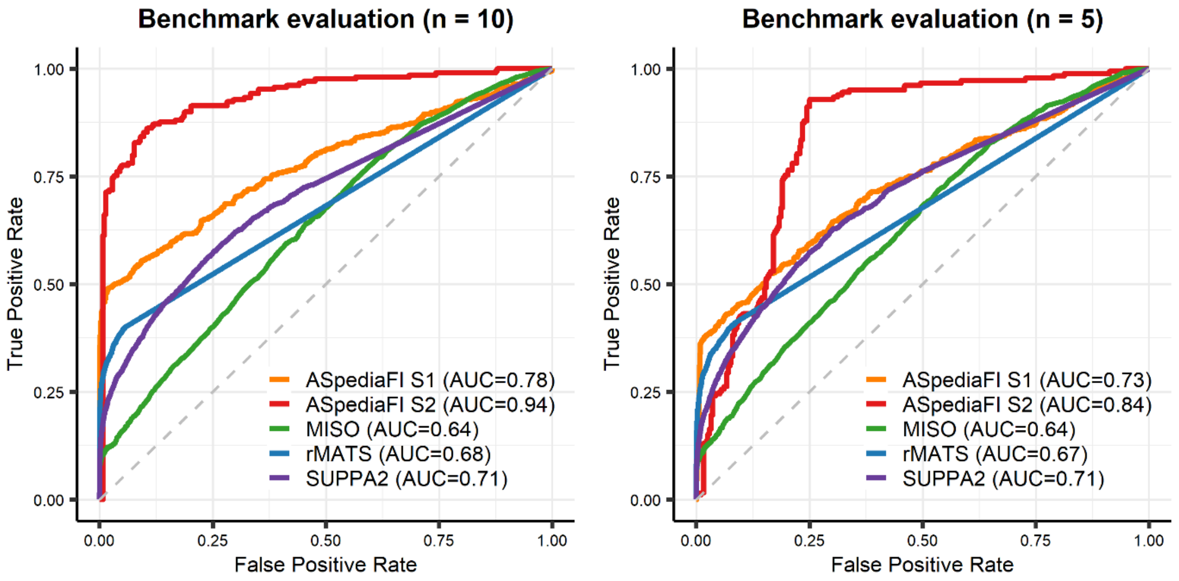

**Figure S4 Precision and recall for HM and HM expansion gene set calculated from a simulated dataset of size 20 replicates per condition.** The same resulting DAS genes from the comparison analysis (Figure 5F) were evaluated. Each bar on the x-axis represents a DAS detection tool with its corresponding color at the legend (Bottom).

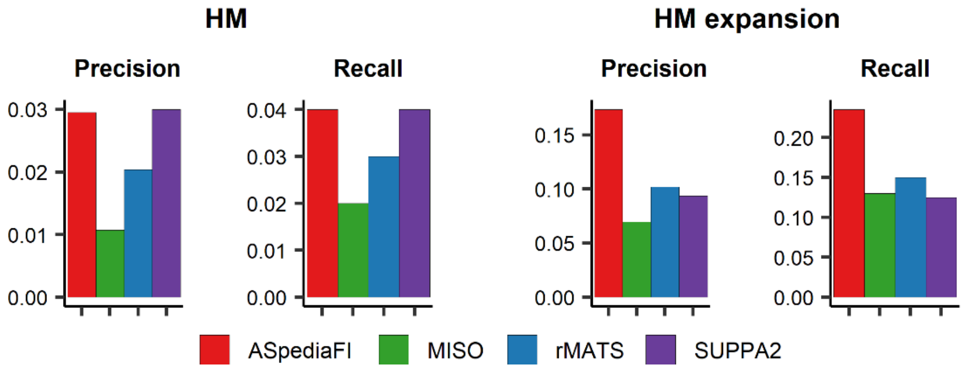
