## Supplementary material for "ASpediaFI: Functional interaction analysis of alternative splicing events": Table S2-S7

**Table S2. Counts of AS events detected by ASpedia, rMATS Cond1 and Cond2.** Counts and percentages of 5 AS types are summarized for three cases and three SF comparisons.

| **Tool** | **AS event type** | **SF3B1** | **SRSF2** | **U2AF1** |
| --- | --- | --- | --- | --- |
| **ASpediaFI** | **A3** | 66 (23.5%) | 60 (22.3%) | 64 (22.5%) |
|  | **A5** | 28 (10.0%) | 53 (19.7%) | 37 (13.0%) |
|  | **SE** | 33 (11.7%) | 50 (18.6%) | 40 (14.0%) |
|  | **RI** | 151 (53.7%) | 102 (37.9%) | 142 (49.8%) |
|  | **MXE** | 3 (1.1%) | 4 (1.5%) | 2 (0.7%) |
|  | **Total** | 281 | 269 | 285 |
| **rMATS**  **Cond1** | **A3** | 102 (17.1%) | 67 (9.3%) | 107 (12.5%) |
|  | **A5** | 37 (6.2%) | 50 (7.0%) | 46 (5.4%) |
|  | **SE** | 241 (40.4%) | 413 (57.4%) | 425 (49.8%) |
|  | **RI** | 137 (23.0%) | 78 (10.8%) | 119 (13.9%) |
|  | **MXE** | 79 (13.3%) | 111 (15.4%) | 157 (18.4%) |
|  | **Total** | 596 | 719 | 854 |
| **rMATS**  **Cond2** | **A3** | 81 (22.1%) | 34 (10.3%) | 54 (17.5%) |
|  | **A5** | 17 (4.6%) | 20 (6.1%) | 12 (3.9%) |
|  | **SE** | 125 (34.1%) | 196 (59.4%) | 168 (54.4%) |
|  | **RI** | 94 (25.6%) | 38 (11.5%) | 39 (12.6%) |
|  | **MXE** | 50 (13.6%) | 42 (12.7%) | 36 (11.7%) |
|  | **Total** | 367 | 330 | 309 |

**Table S3. Exclusively detected AS events overlapping with heme metabolism (HM) expansion gene set.** To compare AS events identified from ours and rMATS Cond2, genes and involved protein domains were listed for gene set to belong HM and novel (expansion set not including HM gene set).

| **ASpediaFI** | | **rMATS Cond2** | |
| --- | --- | --- | --- |
| **HM set**  **(n=5)** | **Novel (HE expansion set; n=17)** | **HM set**  **(n=3)** | **Novel (HE expansion set; n=5)** |
| NARF (Iron only hydrogenase large subunit, C-terminal) | CDC37 (Hsp90 binding) | CAST (Calpain inhibitor) | ANKZF1 (Ankyrin repeats) |
| RAD23A | CHD4 | FBXO7 (PI31 proteasome regulator N-terminal) | CD46 |
| RBM5 (RNA recognition motif) | DDX24 (Helicase conserved C-terminal) | LAMP2 (Lysosome-associated membrane glycoprotein) | CD59 |
| SDCBP (PDZ) | EIF3K |  | MAP3K4 (Protein kinase domain) |
| SNCA (Synuclein) | EWSR1 |  | TUBA1A |
|  | HMGN2 |  |  |
|  | KAT2A |  |  |
|  | MORF4L2 |  |  |
|  | MOV10 (AAA domain) |  |  |
|  | NOL11 |  |  |
|  | NR1H3 (Ligand-binding domain of nuclear hormone receptorf) |  |  |
|  | OGT |  |  |
|  | PIGA (Glycosyl transferases group) |  |  |
|  | PTPN6 (SH2 domain) |  |  |
|  | U2AF1 |  |  |
|  | UBE3A (HECT-domain (ubiquitin-transferase)) |  |  |
|  | VPS51 |  |  |

**Table S4. Counts and percentage of AS events with functional sequence features retrieved from ASpedia database.** The number of AS events involved in protein domain, NMD, PTM, PPI and repeat regions was summarized for case study 1~3. ASpediaFI and other applications’ results were included.

| **Dataset and analysis method** | **Case 1: MDS SF3B1** | | | **Case 2: ESRP1** | | **Case 3: RBFOX1** |
| --- | --- | --- | --- | --- | --- | --- |
|  | **ASpediaFI**  **(n = 281)** | **rMATS1 Cond 1**  **(n = 596)** | **rMATS2 Cond 2**  **(n = 367)** | **ASpediaFI**  **(n = 246)** | **SUPPA**  **(n = 341)** | **ASpediaFI**  **(n = 291)** |
| **Protein domain**  **(Pfam)** | 166 (59.1%) | 301 (50.5%) | 203 (55.3%) | 80 (32.5%) | 113 (33.1%) | 133 (45.7%) |
| **NMD**  **(Ensembl)** | 48 (17.1%) | 105 (17.6%) | 73 (19.9%) | 17 (6.9%) | 12 (3.5%) | 23 (7.9%) |
| **PTM**  **(PhosphoSitePlus)** | 38 (13.5%) | 104 (17.4%) | 68 (18.5%) | 91 (37.0%) | 102 (29.9%) | 52 (17.9%) |
| **PPI**  **(Uniprot)** | 42 (14.9%) | 59 (9.9%) | 41 (11.2%) | 88 (35.8%) | 82 (24.0%) | 62 (21.3%) |
| **Repeat region**  **(UCSC)** | 80 (28.5%) | 243 (40.8%) | 133 (36.2%) | 24 (9.8%) | 52 (15.2%) | 41 (14.1%) |

**Table S5. Exclusively detected AS events overlapping with EMT expansion gene set.** To compare AS events identified from ours and SUPPA2, genes and involved protein domains were listed for gene set to belong EMT and novel.

| **ASpediaFI** | | **SUPPA2 Cond2** | |
| --- | --- | --- | --- |
| **EMT set**  **(n=1)** | **Novel**  **(EMT expansion set; n=23)** | **EMT set**  **(n=7)** | **Novel**  **(EMT expansion set; n=16)** |
| TGFBI | CAMK2G | ABI3BP | ARMCX2 |
|  | CD46 | COL8A2 | ARRB1 (Arrestin, C-terminal domain) |
|  | DNM1L | FLNA (Filamin/ABP280 repeat domain) | CASP8AP2 |
|  | ERBB2IP (PDZ domain) | LAMA2 (Laminin G domain) | CSK |
|  | FGFR2 (Immunoglobulin I-set domain) | MCM7 (MCM N-terminal domain) | DHFR (Dihydrofolate reductase domain) |
|  | ITGA7 | PLOD2 | EIF2S2 |
|  | ITGB4 (Fibronectin type III domain) | SGCD | EMILIN1 |
|  | LTBP3 (Calcium-binding EGF domain) |  | ESP8 |
|  | LTBP4 (Calcium-binding EGF domain) |  | GAB1 |
|  | MAP2K7 |  | MAPK7 |
|  | MAP4K4 |  | PRELP |
|  | MAPT |  | RAC1 (Ras family domain) |
|  | PDLIM7 |  | SIGIRR |
|  | PEX19 |  | SNRK (Protein kinase domain) |
|  | PKN1 |  | TRIO |
|  | PPHLN1 |  | ZNF408 |
|  | PPP1R12A |  |  |
|  | RNF146 |  |  |
|  | SHC2 (Phosphotyrosine interaction domain) |  |  |
|  | SPAG9 |  |  |
|  | TCF12 |  |  |
|  | TCF7L2 (N-terminal CTNNB1 binding domain) |  |  |
|  | TTN (Immunoglobulin I-set domain, Titin Z domain) |  |  |

**Table S6 Summary of performance metrics for DAS analysis tools using MDS dataset with SF3B1 deficiency.** The original results of ASpediaFI (Figure 5A) were compared against the other three tools with another significance conditions applied.

| Method | DAS events | HM | | | | | HM expansion | | | |
| --- | --- | --- | --- | --- | --- | --- | --- | --- | --- | --- |
|  |  | Fisher P (-log10) | Precision | | Recall | F1 score | Fisher P (-log10) | Precision | Recall | F1 score |
| ASpediaFI | 281 | **2.18** | **0.041** | **0.040** | | **0.041** | **2.44** | **0.180** | **0.175** | **0.178** |
| MISO | 300* | 1.72 | 0.030 | 0.030 | | 0.030 | 1.22 | 0.089 | 0.090 | 0.090 |
| rMATS | 300* | 1.47 | 0.028 | 0.030 | | 0.029 | 0.58 | 0.081 | 0.085 | 0.083 |
| SUPPA2 | 300* | 1.27 | 0.029 | 0.035 | | 0.032 | 1.07 | 0.092 | 0.110 | 0.100 |

* Significant DAS events determined as the following: | dPSI | > 0.1 and top 300 events ranked by

*Bayes Factor* for MISO, by *FDR* for rMATS and SUPPA2

**Table S7 Summary of performance metrics for DAS analysis tools using simulated dataset of size 20 replicates per condition.** The original results of ASpediaFI (Figure 5F) were compared against the other three tools with another significance thresholds applied.

| Method | DAS events | HM | | | | HM expansion | | | |
| --- | --- | --- | --- | --- | --- | --- | --- | --- | --- |
|  |  | Fisher P (-log10) | Precision | Recall | F1 score | Fisher P (-log10) | Precision | Recall | F1 score |
| ASpediaFI | 499 | 1.39 | 0.030 | **0.040** | **0.034** | 2.72 | **0.173** | **0.235** | **0.200** |
| MISO | 218* | 0.19 | 0.007 | 0.005 | 0.006 | 0.99 | 0.086 | 0.060 | 0.071 |
| rMATS | 280* | **1.84** | **0.043** | 0.025 | 0.032 | **3.07** | 0.145 | 0.085 | 0.107 |
| SUPPA2 | 242* | 0.64 | 0.039 | 0.020 | 0.026 | 1.02 | 0.106 | 0.055 | 0.072 |

* Significant DAS events determined as the following: | dPSI | > 0.1 and *Bayes Factor* $\geq5$ for MISO, $FDR<5\%$for rMATS and SUPPA2
